# Bayesian Genome-wide TWAS method to leverage both cis- and trans- eQTL information through summary statistics

**DOI:** 10.1101/2020.03.05.979187

**Authors:** Justin M. Luningham, Junyu Chen, Shizhen Tang, Philip L. De Jager, David A. Bennett, Aron S. Buchman, Jingjing Yang

## Abstract

Transcriptome-wide association studies (TWAS) have been widely used to integrate gene expression and genetic data for studying complex traits. Due to the computational burden, existing TWAS methods do not assess distant trans- expression quantitative trait loci (eQTL) that are known to explain important expression variation for most genes. We propose a Bayesian Genome-wide TWAS (BGW-TWAS) method which leverages both cis- and trans-eQTL information for TWAS. Our BGW-TWAS method is based on Bayesian variable selection regression, which not only accounts for cis- and trans- eQTL of the target gene but also enables efficient computation by using summary statistics from standard eQTL analyses. Our simulation studies illustrated that BGW-TWAS achieved higher power compared to existing TWAS methods that do not assess trans-eQTL information. We further applied BWG-TWAS to individual-level GWAS data (N=∼3.3K), which identified significant associations between the genetically regulated gene expression (GReX) of gene *ZC3H12B* and Alzheimer’s dementia (AD) (p-value= 5.42 × 10^−13^), neurofibrillary tangle density (p-value= 1.89 ×10^−6^ ), and global measure of AD pathology (p-value=9.59 × 10^−7^). These associations for gene *ZC3H12B* were completely driven by trans-eQTL. Additionally, the GReX of gene *KCTD12* was found to be significantly associated with *β*-amyloid (p-value= 3.44 ×10 ^−8^) which was driven by both cis- and trans- eQTL. Four of the top driven trans-eQTL of *ZC3H12B* are located within gene *APOC1*, a known major risk gene of AD and blood lipids. Additionally, by applying BGW-TWAS with summary-level GWAS data of AD (N=∼54K), we identified 13 significant genes including known GWAS risk genes *HLA-DRB1* and *APOC1*, as well as *ZC3H12B.*

## Introduction

Although genome-wide association studies (GWAS) have identified thousands of variants associated with complex traits over the past decades^1-5^, most of these associations are located within noncoding regions and the underlying biological mechanisms by which these variants impact a phenotype are unknown^6; 7^. Recent studies have shown that GWAS associations were enriched for regulatory elements such as expression quantitative trait loci (eQTL)^8-10^, suggesting that integrating transcriptomic and genetic data could help identify key molecular mechanisms underlying complex traits.

One such integrative method is transcriptome-wide association study (TWAS)^11-13^, which takes advantage of a reference panel with profiled transcriptomic and genetic data from the same individuals. TWAS first utilizes such reference data to fit an imputation regression model for the expression quantitative trait of a target gene with nearby genotypes (e.g., cis-SNPs within 1MB region of transcription starting site) as predictors, and then examines the gene-based association between the imputed genetically regulated gene expression (GReX) and the phenotype of interest. With fitted gene expression imputation models from reference data, TWAS can be conducted with test samples that have either individual-level or summary-level GWAS data^12-14^. The SNPs with non-zero effect sizes on reference transcriptome in the fitted imputation models are referred to as broad sense “eQTL” in TWAS. Examples of publicly available reference data include the Genotype-Tissue Expression (GTEx) project with transcriptomic data for 54 human tissues^8^, Genetic European Variation in Health and Disease (GEUVADIS) for lymphoblastoid cell lines^15^, and North American Brain Expression Consortium (NABEC) for cortex tissues^16^.

Essentially, TWAS is equivalent to a burden type gene-based test taking “cis-eQTL effect sizes” that are non-zero coefficients of cis-SNPs from the fitted GReX imputation model as their corresponding burden weights^11-13^. By weighting genetic variants using cis-eQTL effect sizes, TWAS assumes the effects of risk genes on the phenotype of interest are potentially mediated through their transcriptome variations. Recent studies of a wide range of complex traits such as schizophrenia, breast cancer, and Alzheimer’s dementia (AD)^17-21^ using TWAS have identified additional risk genes besides known GWAS risk loci, demonstrating that additional significant associations can be identified by TWAS.

However, existing TWAS methods only use genetic data of cis-SNPs of the target gene as predictors to fit the GReX imputation model^11-13^. As shown by recent studies, trans-SNPs (e.g., outside of the 1MB region) of the target gene not only explain a significant amount of variation for most expression quantitative traits, but also often contain significant trans-eQTL that are likely to inform molecular mechanisms^22; 23^. Thus, using both cis- and trans- SNPs is likely to increase the imputation accuracy of GReX and the power of TWAS. Nonetheless, the enormous computational cost required to fit ∼20K GReX imputation models for genome-wide genes and genotypes per tissue type makes the routine use of existing TWAS methods impractical.

We propose a Bayesian Genome-Wide TWAS (BGW-TWAS) method that accounts for both cis- and trans- SNPs based on a Bayesian variable selection regression (BVSR) model^24^ for imputing GReX. Our BGW-TWAS method circumvents the current computational burden impeding TWAS by enabling efficient computation via the scalable EM-MCMC algorithm^25^ and the summary statistics of standard eQTL analyses based on single variant tests. First, we demonstrate the feasibility of this Bayesian approach by simulation studies with varying proportions of true causal cis- and trans- eQTL for expression quantitative traits. We compared BGW-TWAS with several existing TWAS methods including PrediXcan^11^ and TIGAR^13^ that assess only cis-SNPs. Then we applied BGW-TWAS to clinical and postmortem data from older adults with individual-level GWAS data (N=∼3.3K)^26^, to study several clinical and pathological AD related phenotypes including clinical diagnosis of AD, neurofibrillary tangle density, *β*-amyloid load, and a global summary measure of AD pathology. Further, we compared BGW-TWAS with alternative methods by using GWAS summary statistics for AD available from the International Genomics of Alzheimer’s Project (IGAP)^27^ (N=∼54K).

Our simulation studies revealed that BGW-TWAS achieved higher TWAS power by considering both cis- and trans- SNPs when trans-eQTL accounted for a non-negligible proportion of transcriptome variance. Our studies of human AD GWAS datasets identified several risk genes associated with AD phenotypes that were driven by trans-eQTL and thus not identified by alternative methods. The software for implementing BGW-TWAS is available freely on Github.

## Method

### TWAS Procedure

The first step in TWAS is to train an imputation model for profiled gene expression levels using genotype data as predictors on a per-gene basis^11-13^. The general imputation model based on linear regression is given by

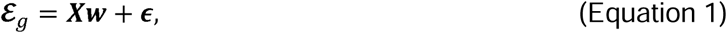

where ***ε***_*g*_ denotes the expression quantitative trait of the target gene, centered and adjusted for non-genetic covariates; ***X*** denotes centered genotype data; ***w*** denotes the corresponding “eQTL” effect sizes for the target gene; and ***ϵ*** is an error term following a 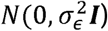 distribution. The intercept term is dropped for centering both response (***ε***_*g*_) and explanatory (***X***) variables. With ***ŵ*** estimated from the training data (i.e., reference data) that have both transcriptomic and genetic data profiled for the same subjects, TWAS will test the association between the phenotype of interest and the imputed GReX obtained from individual-level genotype data 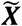 of the test cohort as follows

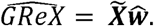

### Bayesian Variable Selection Regression

Existing TWAS methods only consider SNPs within 1MB of the flanking 5’ and 3’ ends (cis-SNPs) in the gene expression imputation model (Equation 1)^11-13^. In order to leverage additional information provided by trans-eQTL that are located outside the 1MB flanking region of the target gene, we utilize the Bayesian Variable Selection Regression (BVSR)^24^ model to account for both cis- and trans- SNPs as follows:

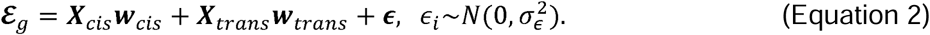

The BVSR model assumes a spike-and-slab prior distribution for *w*_*i*_. That is, the prior on *w*_*i*_ is a mixture distribution of a normal distribution with zero mean and a point-mass density function at 0. In order to model potentially different distributions of the effect sizes for cis- and trans- SNPs, we assume the following respective priors,

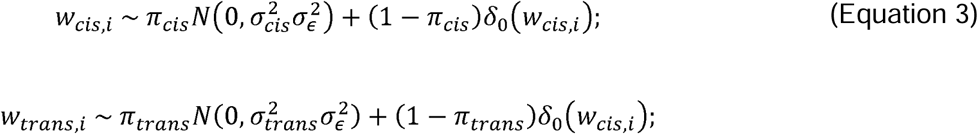

where (*π*_*cis*_, *π*_*trans*_)denote the respective probability that the coefficient is normally distributed, and *δ*_0_(*w*_*i*_) is the point mass density function that takes value 0 when *w*_*i*_ ≠ 0 and 1 when *w*_*i*_ = 0. Further, the following conjugate hyper prior distributions are respectively assumed for the cis- and trans-specific parameters,

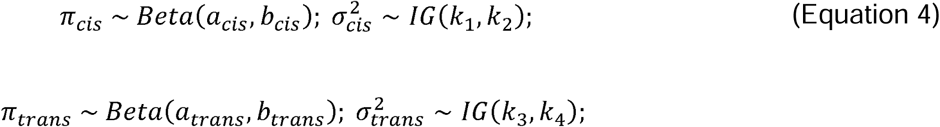

where *IG* indicates the Inverse Gamma distribution and hyper parameters (*a*_*cis*_, *b*_*cis*_, *a*_*trans*_, *b*_*trans*_, *k*_1_, *k*_2_, *k*_3_, *k*_4_) will be chosen to enable non-informative hyper prior distributions (see Supplemental Methods for model details).

To facilitate computation, a latent indicator *γ*_*i*_ is assumed such that *w*_*i*_ =0 if *γ*_*i*_ = 0, and *w*_*i*_ follows a normal distribution if *γ*_*i*_ = 1. Then the expected value of this indicator, E[*γ*_*i*_], represents the posterior probability (*PP*_*i*_) for each individual SNP to have a non-zero effect size (i.e., to be an eQTL of the target gene)^24^. Moreover, we propose a Bayesian approach to estimate GReX for test samples that can account for the uncertainty for each SNP to be an eQTL:

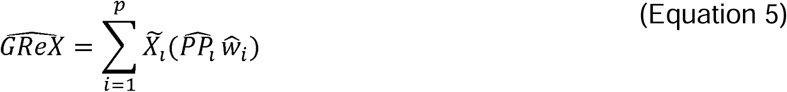

where 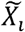 represents the genotype data of variant *i* for test samples and 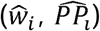 denote the estimate of effect size and posterior probability (PP) of having a non-zero effect size from the BVSR model (Equations 2-4) (see Supplemental Methods for detailed Bayesian inference procedure). This Bayesian GReX estimate can then be used to conduct TWAS with individual-level GWAS data by testing the association between the imputed GReX and the phenotype of interest.

### BGW-TWAS with Summary-level GWAS Data

With summary-level GWAS data that were generated by single variant tests, we employed the S-PrediXcan^14^ approach to obtain a burden TWAS Z-score test statistic, including not only cis-but also trans-eQTL in the test. Let 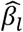 denote the SNP effect size of SNP *l* from GWAS, 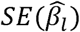 denote the standard error of 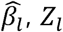 denote the Z-score statistic value by single variant test, 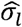 denote the estimated standard deviation of the genotype data of SNP *l* from reference panel, and 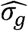 denote the estimated standard deviation of the imputed expression of gene ***g*** from reference panel. The burden TWAS Z-score test statistic for gene ***g*** is given by

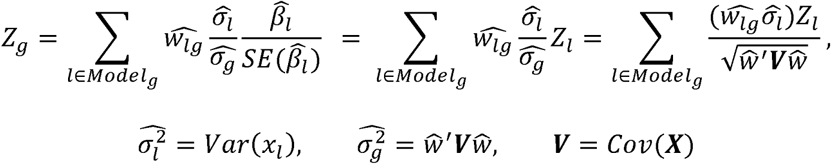

where 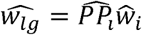 is the product of posterior probability for SNP *l* to have non-zero eQTL effect size from the BVSR model (Equation 2). Here, ***X*** denotes the genotype matrix of analyzed SNPs from reference panels of the same ethnicity and ***V*** denotes the corresponding genotype covariance matrix.

### Efficient Computation Techniques

In theory, the estimates of eQTL effect sizes and corresponding posterior probabilities 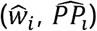 can be obtained by using a standard Markov Chain Monte Carlo (MCMC)^28^ algorithm. However, in practice, the computation burden for modeling genome-wide genotype data is nearly impossible because of enormous required memory capacity and slow convergence rate for MCMC. To circumvent these practical limitations, we employ several techniques to enable computational efficiency such that BGW-TWAS method can be deployed to leverage both cis- and trans- eQTL information in practice. In particular, we adapt a previously developed scalable expectation-maximization Markov Chain Monte Carlo (EM-MCMC) algorithm^25^. Unlike the original EM-MCMC algorithm requiring individual-level GWAS data, we can reduce up to 90% of the computation time by adapting the EM-MCMC algorithm to utilize only summary statistics, including the pre-calculated linkage disequilibrium (LD) coefficients and score statistics from standard eQTL analyses by single variant tests. Additionally, we prune genome-wide genotypes into a subset of genome regions that are approximately independent and contain either at least one cis-SNP or one trans-SNP with p-value <1 × 10 ^−5^ by standard eQTL analyses.

### Simulation Study Design

We conducted simulation studies to validate the performance of our proposed BGW-TWAS method through comparing with the alternative existing methods, e.g., PrediXcan, TIGAR, as well as BVSR using only cis-eQTL. To mimic real studies, we used real genotype data from the ROS/MAP study to simulate gene expression and phenotype data. We took 499 samples as our training data and 1,209 samples as our test data. GReX imputation models were fitted using the training data where “eQTL” effect sizes and the corresponding posterior probabilities were estimated. Given these fitted GReX imputation models, GReX data were imputed for follow-up TWAS with the test data.

We arbitrarily selected five approximately independent genome blocks, including one “cis-” and four “trans-” genotype blocks (variants were filtered with minor allele frequency (MAF) > 5% and Hardy-Weinberg p-value > 10^−5^). With genotype matrix ***X***_*****g*****_ of the randomly selected causal eQTL, we generated effect-sizes *w*_*i*_ to target a selected gene expression heritability 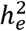 and that all causal eQTL explain equal expression heritability. Gene expression levels were generated by ***ε***_***g***_ = ***X***_***g***_***w***+ *ϵ*_*e*_, with 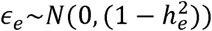. Then we simulated phenotypes by ***Y***= *β****ε***_***g***_ + *ϵ*_*p*_, where *β* was selected with respect to a selected phenotype heritability 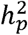 and 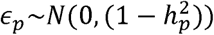.

To mimic the complex genomic architecture of gene expression in practice, we considered two scenarios, one with 5 true causal eQTL representing the scenario with relatively large effect sizes and the other one with 22 true causal eQTL representing the scenario with relatively small effect sizes. For the scenario with 5 true causal eQTL, we considered three sub-scenarios with respect to how these true causal eQTL distributed over considered genome blocks: i) all causal eQTL are from the cis-block; ii) two causal eQTL are from the cis-block explaining 70% of the specified 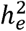 while the other three causal eQTL are from the trans-blocks explaining the other 30% of 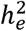; iii) all causal QTL are from the trans-block. Similarly, For the scenario with 22 true causal eQTL, we considered three scenarios where 30%, 50%, and 70% of the causal eQTL were from *cis-* genome blocks. We also varied the total expression trait heritability and phenotype heritability in both scenarios, i.e., 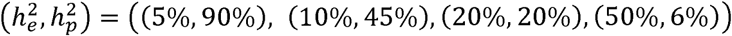 for the scenario with 5 true causal eQTL and 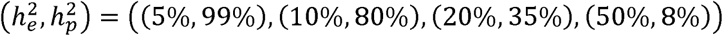 for the scenario with 22 true causal eQTL. Here, different levels of phenotype heritability were arbitrarily selected to achieve similar levels of TWAS power across all scenarios.

In each simulation, with training data, we first fitted GReX imputation models by BVSR (BGW-TWAS) with both *cis*- and *trans*- genome blocks, as well as by Elastic-Net (PrediXcan) and nonparametric Bayesian Dirichlet process regression (TIGAR) with only *cis*- genome block. Then we conducted TWAS with imputed *GReX* by respective method. We also compared BGW-TWAS with using only cis-eQTL estimates from the same BVSR model. The performance was compared in terms of *R*^2^ of the imputed GReX and TWAS power in test samples. Test *R*^2^ was calculated as the squared correlation between imputed GReX and simulated gene expression values of the test samples. TWAS power was calculated as the proportion of 1,000 repeated simulations of each scenario with *p*-value < 2.5 × 10 ^−6^ (genome-wide significance threshold for gene-based association studies).

### ROS/MAP and Mayo Clinic GWAS data of AD

Following simulation studies, we applied BGW-TWAS method to individual-level genomic and AD related phenotype data from older adults available from several studies. We used transcriptomic data, GWAS data, clinical diagnosis of AD and postmortem indices of AD pathology from the Religious Orders Study (ROS) and Rush Memory and Aging Project (MAP) ^29-31^ and GWAS data from the Mayo Clinic Alzheimer’s Disease Genetics Studies (MCADGS)^32-34^. All participants from ROS/MAP sign an informed consent, an Anatomic Gift Act, and a consent for their data to be deposited in the Rush Alzheimer’s Disease Center (RADC) repository. ROS/MAP studies were approved by the Institutional Review Board of Rush University Medical Center, Chicago, IL. MCADGS contains samples from two clinical AD Case-Control series (Mayo Clinic Jacksonville and Mayo Clinic Rochester) as well as a neuropathological series of autopsy-confirmed subjects from the Mayo Clinic Brain Bank.

Microarray genotype data generated for 2,093 European-decent subjects from ROS/MAP^35^ and 2,099 European-decent subjects from MCADGS were further imputed to the 1000 Genome Project Phase 3^36^ in our analysis^37^.

Post-mortem brain samples from the dorsal lateral prefrontal cortex from ∼30% of these ROS/MAP participants with assayed genotype data were profiled for transcriptomic data by next-generation RNA-seqencing^38^. These data were used as reference data to train GReX prediction models in this study. We conducted TWAS for both clinical and pathological AD phenotypes. The clinical diagnosis of late-onsite Alzheimer’s dementia was available for both ROS/MAP and MCADGS. Postmortem pathology indices of AD were only available for ROS/MAP and included PHFtau tangle density, *β* -amyloid load, and a global measure of AD pathology based on measures of neuritic and diffuse plaques and neurofibrillary tangles^29-31^. Additional details about the ongoing ROS/MAP cohort studies and how postmortem indices of tangles and *β* -amyloid load were quantified are included in prior publications^29-31^ and summarized in the Supplemental Text.

## Results

### Simulation Results

For the scenario with 5 true causal eQTL and various expression heritability, our simulation studies showed that BGW-TWAS obtained the highest test *R*^2^ for GReX and TWAS power than PrediXcan and TIGAR when any portion of the true causal eQTL are distributed over trans- genome blocks (Figure 1 (A, B)). This is because BGW-TWAS leverages both cis- and trans- eQTL information while the alternative methods fail to account for trans- eQTL. Especially, when all true causal eQTL are from trans- genome regions, the alternative methods barely have any power to identify the TWAS association with nearly zero test *R*^2^. As expected, BGW-TWAS and PrediXcan performed comparably when all causal eQTL were from the cis- genome block, while TIGAR performed slightly worse with sparse true causal eQTL (Figure 1A).

**Figure 1.**
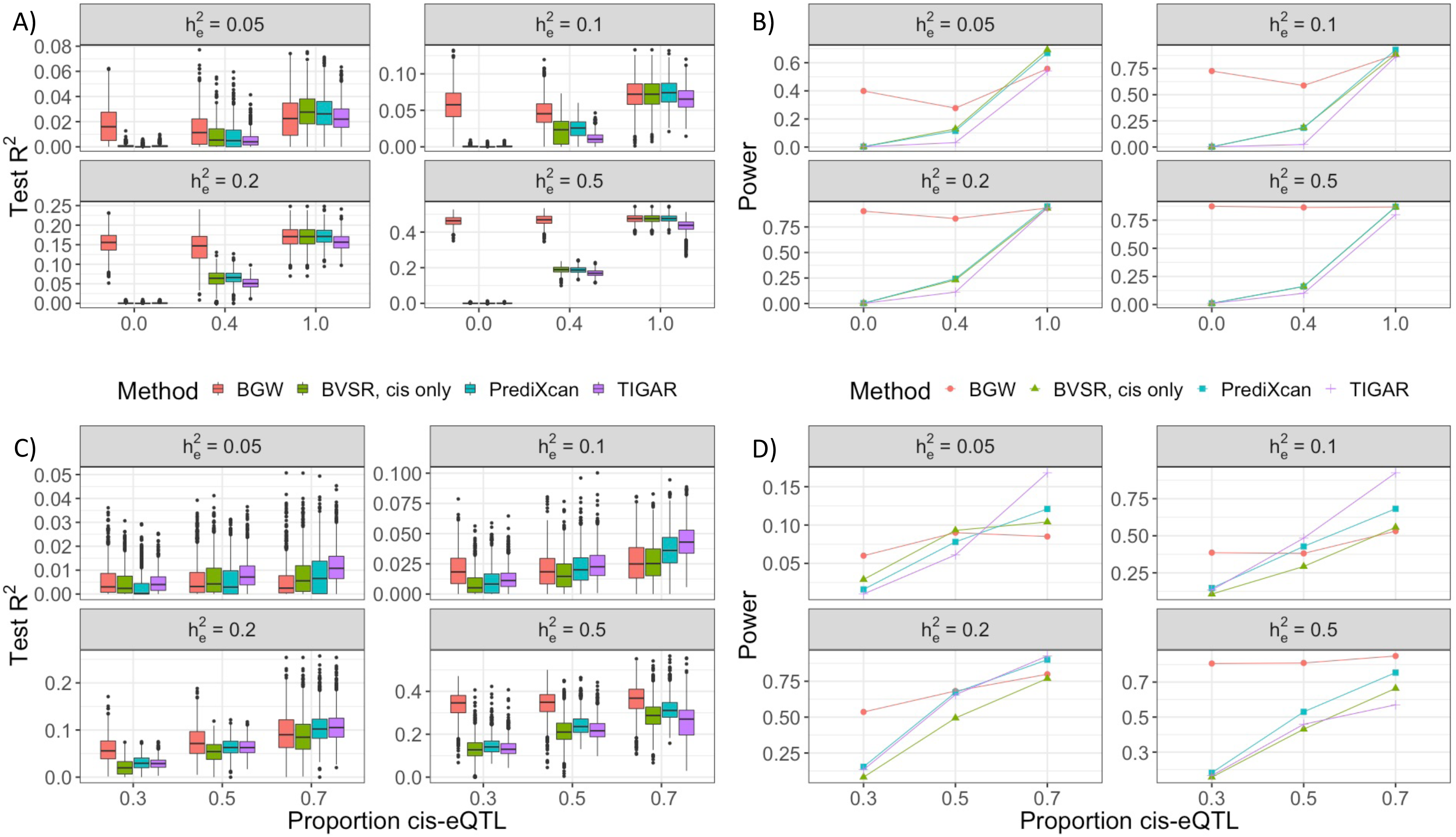
Simulation TWAS Studies Comparing BGW-TWAS, BVSR with cis-eQTL only, PrediXcan and TIGAR Methods. Simulation studies used various gene expression heritability 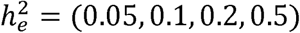 and various true causal cis-eQTL proportions. (A, B) Test R2 and TWAS power comparison with 5 true causal eQTL. BGW-TWAS was found to out-perform the alternative methods when a non-negligible proportion of true causal eQTL were from trans-genome regions. (C, D) Test R2 and TWAS power comparison with 22 true causal eQTL. BGW-TWAS was found to out-perform alternative method when >50% of true causal eQTL were from trans-genome regions and 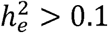.

For the scenario with 22 mixed cis- and trans- eQTL, the performance comparison became more complicated with respect to various true expression heritability levels (Figure 1 (C, D)). Particularly, when 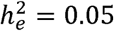, all methods had difficulties accurately estimating eQTL effect sizes and resulted in nearly zero test *R*^2^. As expression heritability increased, the advantage of modeling both cis- and trans- genotype data by BGW-TWAS arisen and led to higher test *R*^2^ and TWAS power. When 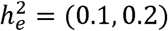 and 70% of the true causal eQTL were cis-, BGW-TWAS was less effective than PrediXcan and TIGAR while TIGAR achieved the best performance. This is likely due to the fact that the nonparametric Bayesian Dirichlet process regression model used by TIGAR is preferred when true causal eQTL manifest relatively small effect sizes, which is consistent with previous findings^13^.

In contrast, when true causal eQTL signals have relatively large effect sizes and are distributed outside the cis-region of the target gene, BGW-TWAS method is preferred due to the improved accuracy for GReX prediction by leveraging trans- SNP data. By comparing with using only BVSR estimates of cis-eQTL, we showed that a significant proportion of transcriptome variation due to trans-eQTL was missed and the follow-up TWAS was underpowered.

### TWAS of AD Related Phenotypes with Individual-level GWAS Data

Next, we applied BGW-TWAS to conduct TWAS using the individual-level GWAS data from ROS/MAP^26; 31^ and MCADGS^32^. First, we trained the BVSR GReX imputation models using samples (N=499) from the ROS/MAP cohort that contained both profiled genotype data and transcriptomic data obtained from the dorsal lateral prefrontal cortex. All expression quantitative traits were normalized and corrected for age at death, sex, postmortem interval (PMI), study (ROS or MAP), batch effects, RNA integrity number (RIN), top three principal components derived from genome-wide genotype data, and cell type proportions (oligodendrocytes, astrocytes, microglia, neurons). The cell type proportions were derived by using CIBERSORT pipeline^39^ with single-cell RNA-seq transcriptome profiles from human brain tissues as the reference^40^ to de-convolute bulk RNA-seq data^41^.

When we applied BGW-TWAS, we obtained GReX imputation models for 14,156 genes, compared to respective 6,011 genes and 14,214 genes by PrediXcan and TIGAR that have at least one cis-eQTL with nonzero effect size on expression quantitative trait. Across the 6,011 genes with GReX imputation models by PrediXcan, our BGW-TWAS approach had a smaller train *R*^2^ (squared correlation between fitted and profiled gene expression values) value for expression quantitative traits for only 855 genes (Figure S1A). While TIGAR and BGW-TWAS yielded a similarly number of GReX imputation models, BGW method is expected to result in higher imputation accuracy when trans-eQTL play an important role in affecting gene expression levels as shown by our simulation results. Of 13,142 genes that had imputation models fitted by both TIGAR and BGW-TWAS, BGW-TWAS had smaller train *R*^2^ for only 3,304 genes. That is, BGW-TWAS would be preferred for genes that have sparse eQTL, especially trans-eQTL, while TIGAR would be preferred for genes that have less sparse eQTL that are mostly cis-eQTL (Figure S 1B).

We imputed Bayesian GReX values for all remaining individuals with genotype data in ROS/MAP and MCADGS by using Equation 4. We then conducted TWAS by testing the association between the standardized GReX values (with unit variance) and both clinical and pathological AD phenotypes. TWAS for these phenotypes controlled for age at death, sex, smoking, ROS or MAP study, education level, and top three principal components derived from genome-wide genotype data.

For the dichotomous phenotype of clinical diagnosis of AD, the case/control status was determined by different rules and the available confounding variables were different for ROS/MAP and MCADGS. Cognitive status at death for individuals from the ROS/MAP cohort is based on the review of all longitudinal clinical data available at the time of death blinded to all pathologic data. Individuals were classified as having no cognitive impairment, mild cognitive impairment or AD. In this study, samples from individuals with AD were taken as cases and samples from individuals without dementia i.e., either with no cognitive impairment or mild cognitive impairment were taken as controls. For the MCADGS samples, cases were determined for samples with a medical history of late onsite AD diagnosis, and available confounding variables only included age, sex, and top three principal components derived from GWAS data. Therefore, we meta-analyzed these two cohorts for AD clinical diagnosis by applying the inverse-variance weighting method^42^ to summary statistics obtained by TWAS per cohort. We compared the meta-TWAS results obtained with BGW-TWAS to alternative TWAS methods.

BGW-TWAS identified gene *ZC3H12B* (located on chromosome X) whose GReX values were associated with AD with effect size *β* = 0.265, p-value 5.42 × 10^−13^, and FDR = 3.07 ×10 ^−8^ (Figure 2A; Table 1). Both within-cohort TWAS obtained positive effect sizes (β= 0.22,0.29) for ROS/MAP and MCADGS, with respective p-value =2 × 10 ^−4^4.12 × 10 ^−10^ On the other hand, this gene was not identified by either PrediXcan or TIGAR because the association of this gene is completely drive by trans-eQTL (Figures S2 and S4).

**Table 1.** Significant risk genes identified by BGW-TWAS using individual-level GWAS data from ROS/MAP and MCADGS cohorts.

| Gene | CHR | Position | Train $R^2$ | p-value | Effect size (SD) | Phenotype |
| --- | --- | --- | --- | --- | --- | --- |
| *ZC3H12B* | X | 64,708,614 | 0.24 | $5.42 \times 10^{-13}$ | 0.265 (0.037) | AD |
| *ZC3H12B* | X | 64,708,614 | 0.24 | $9.59 \times 10^{-7}$ | 0.142 (0.029) | Global AD pathology |
| *ZC3H12B* | X | 64,708,614 | 0.24 | $1.89 \times 10^{-6}$ | 0.138 (0.029) | Tangles |
| *KCTD12* | 13 | 77,454,311 | 0.09 | $3.44 \times 10^{-8}$ | 0.143 (0.026) | β-Amyloid |

**Figure 2.**
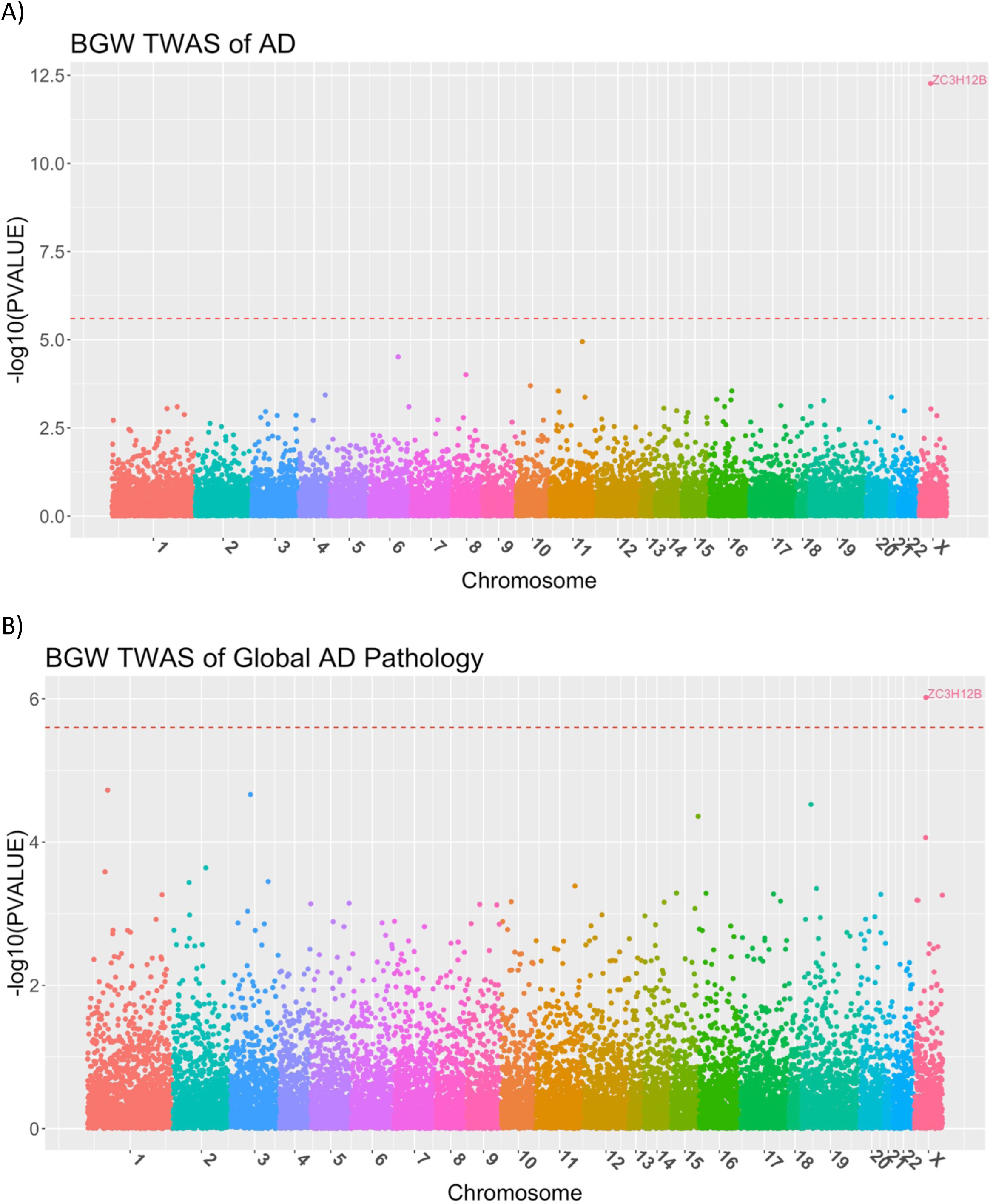
Manhattan plots of BGW-TWAS results of AD clinical diagnosis (A) and global AD pathology (B). Red lines denote genome-wide significant threshold (2.5 × 10^−6^) for gene-based association studies. Gene ZC3H12B was found to be significantly associated with both AD clinical diagnosis and global AD pathology.

TWAS of pathological AD phenotypes were restricted to ROS/MAP from whom postmortem AD indices were collected. TWAS were conducted for individuals with GWAS genotype data and AD pathology indices –– tangles (N=1,121), *β*-amyloid (N=1,114), and global AD pathology (1,139). These results are shown in the Manhattan plots in Figures 2B, 3(A, B) and Figures S2-S3 and S5-S7.

**Figure 3.**
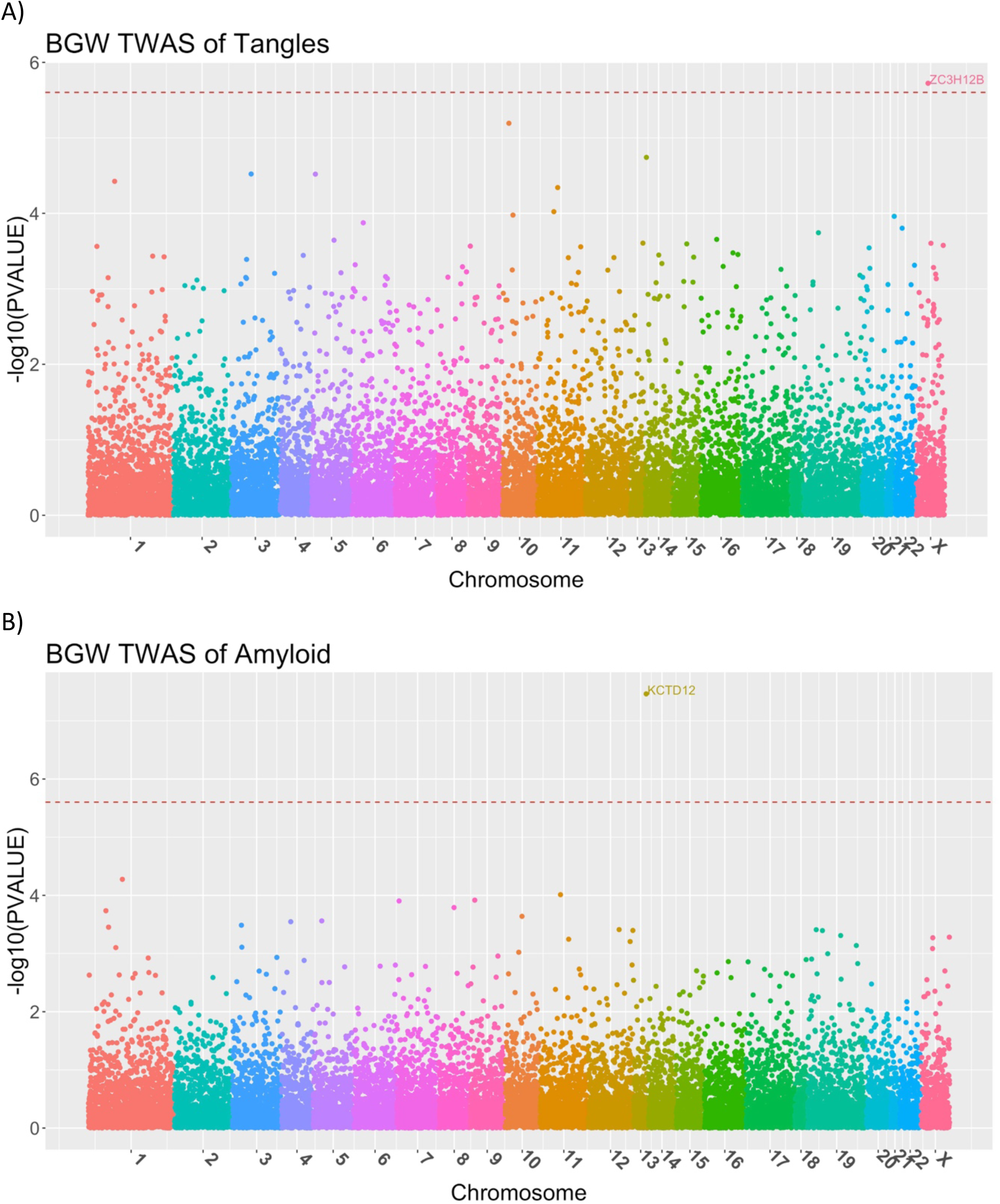
Manhattan plots of BGW-TWAS results of neurofibrillary tangle density (A) and -amyloid load (B). Red lines denote genome-wide significant threshold (2.5 × 10^−6^) for gene-based association studies. Gene ZC3H12B was found to be significantly associated with neurofibrillary tangle density. Gene KCTD12 was found to be significantly associated with and β-amyloid load.

Using BGW-TWAS, *ZC3H12B* was identified to be associated with global AD pathology with p-value = 9.59 × 10 ^−7^ (Figure 2B; Table 1) as well as neurofibrillary tangle density with p-value = 1.89 × 10^−6^ (Figure 3A; Table 1). Gene *KCTD12* located on chromosome 12 was identified to be significantly associated with *β*-amyloid load with p-value = 3.44 × 10 ^−8^ (Figure 3B; Table 1).

We show the BVSR posterior probabilities for considered SNPs to be eQTL for *ZC3H12B* and *KCTD12* in Figure 4, and the standard eQTL analyses results for these two genes in Figure S4. These data suggest that that the association between the imputed GReX values of *ZC3H12B* and AD phenotypes is completely driven by trans-eQTL, while the association between the GReX values of *KCTD12* and *β*-amyloid load is driven by both cis- and trans- eQTL. Four of the top driven trans-eQTL (*rs12721051, rs4420638, rs56131196, rs157592;* Table 2) for *ZC3H12B* are located in gene *APOC1* that is a known risk gene for AD^43^ and blood lipids^44-46^, which is <12KB away from the well-known AD risk gene *APOE*^47^. Particularly, *rs12721051* located in the 3’ UTR region of *APOC1* was identified as a GWAS signal of total cholesterol levels^46^; *rs4420638* located in the downstream of *APOC1* is in linkage disequilibrium (LD) with the *APOE-E4* allele (*rs429358*) and was identified to be a GWAS signal of various blood lipids measurements (i.e., low density lipoprotein cholesterol measurement, C-reactive protein measurement, triglyceride measurement, and total cholesterol measurement)^44^ and AD^48^; *rs56131196* located in the downstream and *rs157592* located in the regulatory region of *APOC1* were identified as GWAS signals of AD and independent of *APOE-E4*^*49*^. Additionally, gene *ZC3H12B* was found to regulate pro-inflammatory activation of macrophages^50^ and has higher expression in brain, spinal cord and thymus tissue types compared to other tissues^51^. These results showed that the effects of these known GWAS signals (*rs4420638, rs56131196, rs157592*) of AD could be mediated through the gene expression levels of *ZC3H12B*.

**Table 2.**
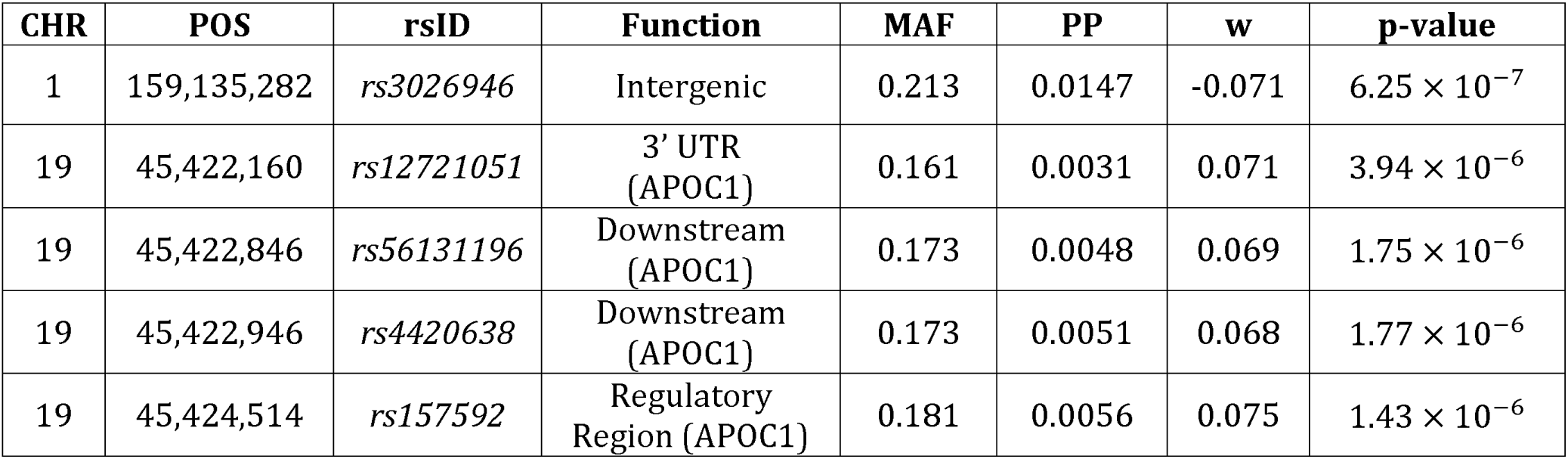
Trans-SNPs with top five posterior probability (PP) > 0.003 of having non-zero eQTL effect sizes for gene ZC3H12B.

**Figure 4.**
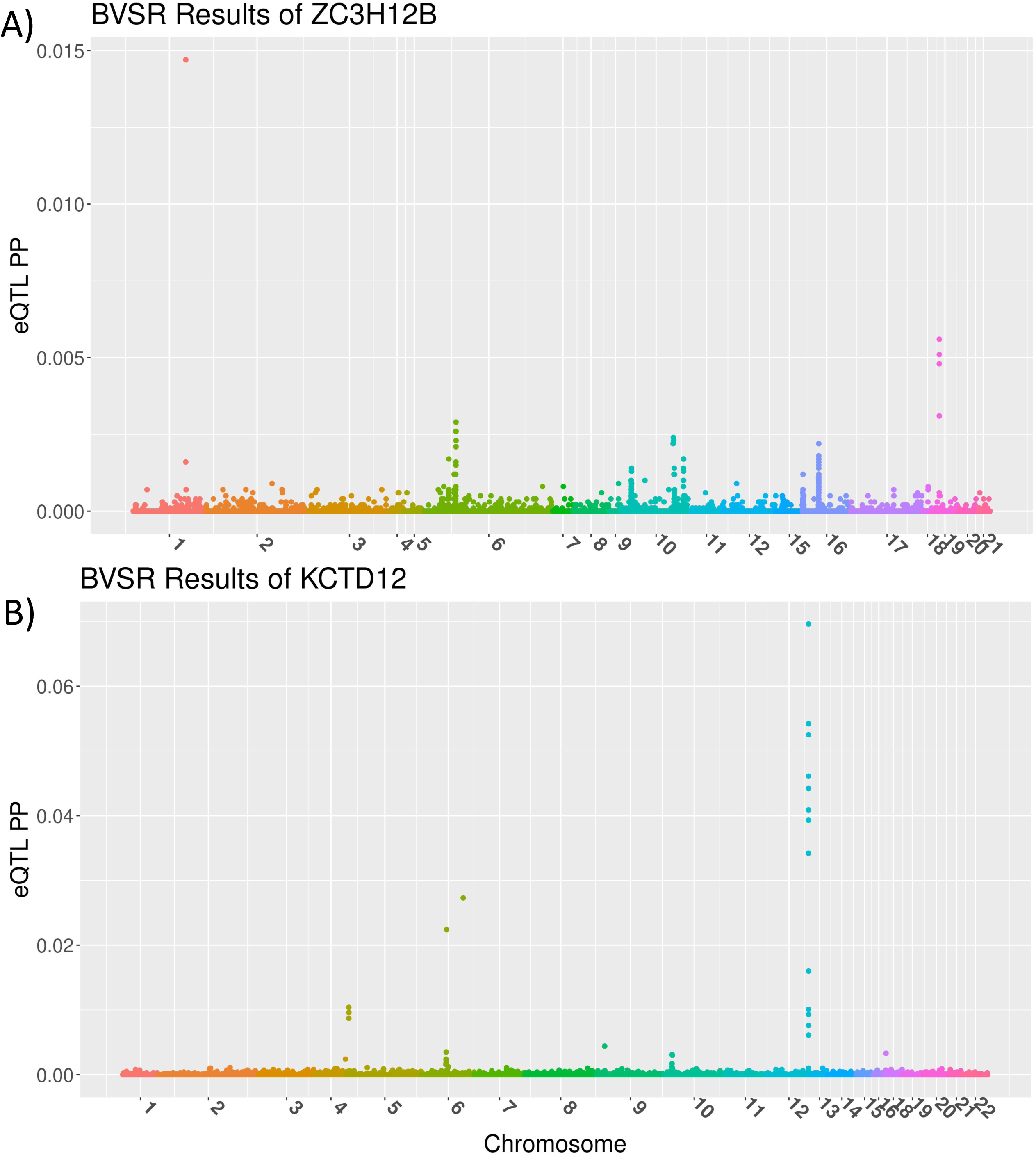
BVSR posterior probabilities (PP) of having non-zero eQTL effect sizes for analyzed cis- and trans- SNPs, with target genes ZC3H12B (A) and KCTD12 (B). Gene ZC3H12B located on chromosome X has top trans-eQTL from chromosomes 1, 6, and 19, where all eQTL are of trans-eQTL. Gene KCTD12 located on chromosome 12 has top cis-eQTL from chromosome 12 and trans-eQTL from chromosomes 4 and 6.

### TWAS of AD with Summary-level GWAS Data

To validate our findings using individual-level GWAS data from ROS/MAP and MCADGS, we conducted TWAS of AD using the publicly available IGAP GWAS summary statistics^27^. Specifically, we used the GWAS summary statistics that were generated by meta-analysis of four consortia (∼17K cases and ∼37K controls, Europeans): the Alzheimer’s Disease Genetic Consortium (ADGC), the Cohorts for Heart and Aging Research in Genomic Epidemiology (CHARGE) Consortium, the European Alzheimer’s Disease Initiative (EADI) and the Genetic and Environmental Risk in Alzheimer’s Disease (GERAD) Consortium.

BGW-TWAS identified 13 significant genes located in chromosome 3, 6, 7, 10, 11, 19, and X, including known GWAS risk genes *HLA-DRB1*^*52*^ and *APOC1*, and gene *ZC3H12B* that was identified using individual-level GWAS data from ROS/MAP and MCADGS (Table 3). Moreover, 7 of these genes (including *HLA-DRB1* and *APOC1*) were also identified when we only considered cis-eQTL estimates by BVSR in TWAS. Gene *CEACAM19* near the well-known GWAS risk gene *APOE* was also identified by S-PrediXcan and TIGAR. Known GWAS risk gene *HLA-DRB1*^*52*^ was also identified by S-PrediXcan (Table 3; Tables S1-S3).

**Table 3.** Significant genes identified by BGW-TWAS using IGAP GWAS summary statistics of AD. TWAS p-values by alternative methods, i.e., using BVSR cis-eQTL estimates only, PrediXcan, and TIGAR are also listed. P-VALUEs for genes that were missed by TWAS were indicated as “-”. Significant gene *ZC3H12B* that was identified by BGW-TWAS using individual-level GWAS data from ROS/MAP and MCADGS was also identified by BGW-TWAS using IGAP summary-level GWAS statistics. Gene *CEACAM19* from chromosome 19 was identified by all TWAS methods, and gene *HLA-DRB1* from chromosome 6 is a known GWAS risk locus.

| Gene | CHR | Position | TWAS P-VALUE |  |  |  |
| --- | --- | --- | --- | --- | --- | --- |
|  |  |  | BGW-TWAS | BVSR cis-eQTL | PrediXcan | TIGAR |
| *GPX1*<sup>a</sup> | 3 | 49,394,608 | $2.45 \times 10^{-98}$ | $2.45 \times 10^{-98}$ | - | $3.15 \times 10^{-1}$ |
| *FAM86DP* | 3 | 75,484,261 | $1.55 \times 10^{-13}$ | $4.81 \times 10^{-1}$ | $5.38 \times 10^{-1}$ | $9.63 \times 10^{-1}$ |
| *BTN3A2*<sup>a</sup> | 6 | 26,378,546 | $1.59 \times 10^{-26}$ | $1.56 \times 10^{-26}$ | $3.17 \times 10^{-1}$ | $5.04 \times 10^{-1}$ |
| *ZNF192*<sup>a</sup> | 6 | 28,124,089 | $1.26 \times 10^{-32}$ | $1.25 \times 10^{-32}$ | $8.56 \times 10^{-2}$ | $2.07 \times 10^{-1}$ |
| *AL022393.7*<sup>a</sup> | 6 | 28,144,452 | $3.25 \times 10^{-178}$ | $2.24 \times 10^{-178}$ | $1.50 \times 10^{-1}$ | $8.36 \times 10^{-2}$ |
| ***HLA-DRB1***<sup>ab</sup> | **6** | **32,557,625** | **$1.02 \times 10^{-12}$** | **$8.99 \times 10^{-13}$** | **$2.06 \times 10^{-6}$** | - |
| *AEBP1* | 7 | 44,154,161 | $5.55 \times 10^{-220}$ | $8.62 \times 10^{-1}$ | $6.69 \times 10^{-1}$ | $4.19 \times 10^{-1}$ |
| *BUB3* | 10 | 124,924,886 | $6.64 \times 10^{-18}$ | $1.05 \times 10^{-2}$ | - | $4.76 \times 10^{-1}$ |
| *FBXO3* | 11 | 33,796,089 | $1.48 \times 10^{-9}$ | $6.88 \times 10^{-1}$ | - | $1.13 \times 10^{-1}$ |
| ***CEACAM19***<sup>abc</sup> | **19** | **45,187,631** | **$4.7 \times 10^{-13}$** | **$2.54 \times 10^{-13}$** | **$3.60 \times 10^{-12}$** | **$2.83 \times 10^{-16}$** |
| *APOC1*<sup>a</sup> | 19 | 45,422,606 | $8.9 \times 10^{-11}$ | $1.11 \times 10^{-10}$ | $3.18 \times 10^{-6}$ | $7.2 \times 10^{-3}$ |
| ***ZC3H12B*** | **X** | **64,727,767** | **$2.08 \times 10^{-37}$** | - | - | - |
| *CXorf56* | X | 118,699,397 | $6.02 \times 10^{-07}$ | - | - | - |

a. Genes that were also identified as significant by using BVSR cis-eQTL estimates.
b. Genes that were also identified by PrediXcan.
c. Genes that were also identified by TIGAR.

Our results showed that by using BVSR estimates of cis- and trans- eQTL (BGW-TWAS), most independent risk loci were identified including loci driven by trans-eQTL. For those significant genes driven mainly by cis-eQTL, TWAS using BVSR estimates of cis-eQTL still identified more independent significant risk loci (distributed over chromosomes 2, 3, 6, 11, and 19) than S-PrediXcan and TIGAR, including all 4 significant genes (*HLA-DRB1, SLC39A13, PVR, CEACAM19*) identified by S-PrediXcan and 4 out of 21 significant genes (*ZNF227, ZFP112, PVR, CEACAM19*) identified by TIGAR (Tables S1-3S). Although TIGAR identified most significant TWAS genes (21), these genes are from chromosomes 11 and 19, which are likely to be driven by the same cis-eQTL from two independent loci.

These TWAS results using summary-level GWAS data with a much larger sample size validated our findings obtained with BGW-TWAS using individual-level GWAS data from ROS/MAP and MCADGS.

### Insights about eQTL Genetic Architecture

In addition to imputing Bayesian GReX values (Equation 5), the posterior probabilities of having non-zero eQTL effect sizes estimated by BVSR also provide insights into the genetic architecture of eQTL, especially about how potential eQTL are distributed across the genome. Note that the posterior probability obtained from the BVSR model (Equations 2-4) is essentially the expected probability for a SNP to be an eQTL. Therefore, the sum of posterior probabilities of having non-zero eQTL effect sizes represents the expected number of eQTL.

From our simulation studies, we observed that the expected proportions of cis-eQTL were consistent with the true proportions of causal cis-eQTL. The expected number of eQTL obtained across simulation scenarios is presented in Table 4, where two out of five (40%) causal eQTL and 11 out of 22 (50%) causal eQTL are from cis- genome regions. We can see that, with higher true expression heritability, the expected number of eQTL is closer to the true number of causal eQTL. We can also see that the expected number of eQTL is more accurate for the scenario with 5 true causal eQTL than with 22 true causal eQTL, which is due to the fact that BVSR model prefers relatively larger eQTL effect sizes. These simulation results demonstrated the validity of our BGW-TWAS method based on the BVSR model as well as the usefulness of the sum of posterior probabilities of having non-zero eQTL effect sizes.

**Table 4.**
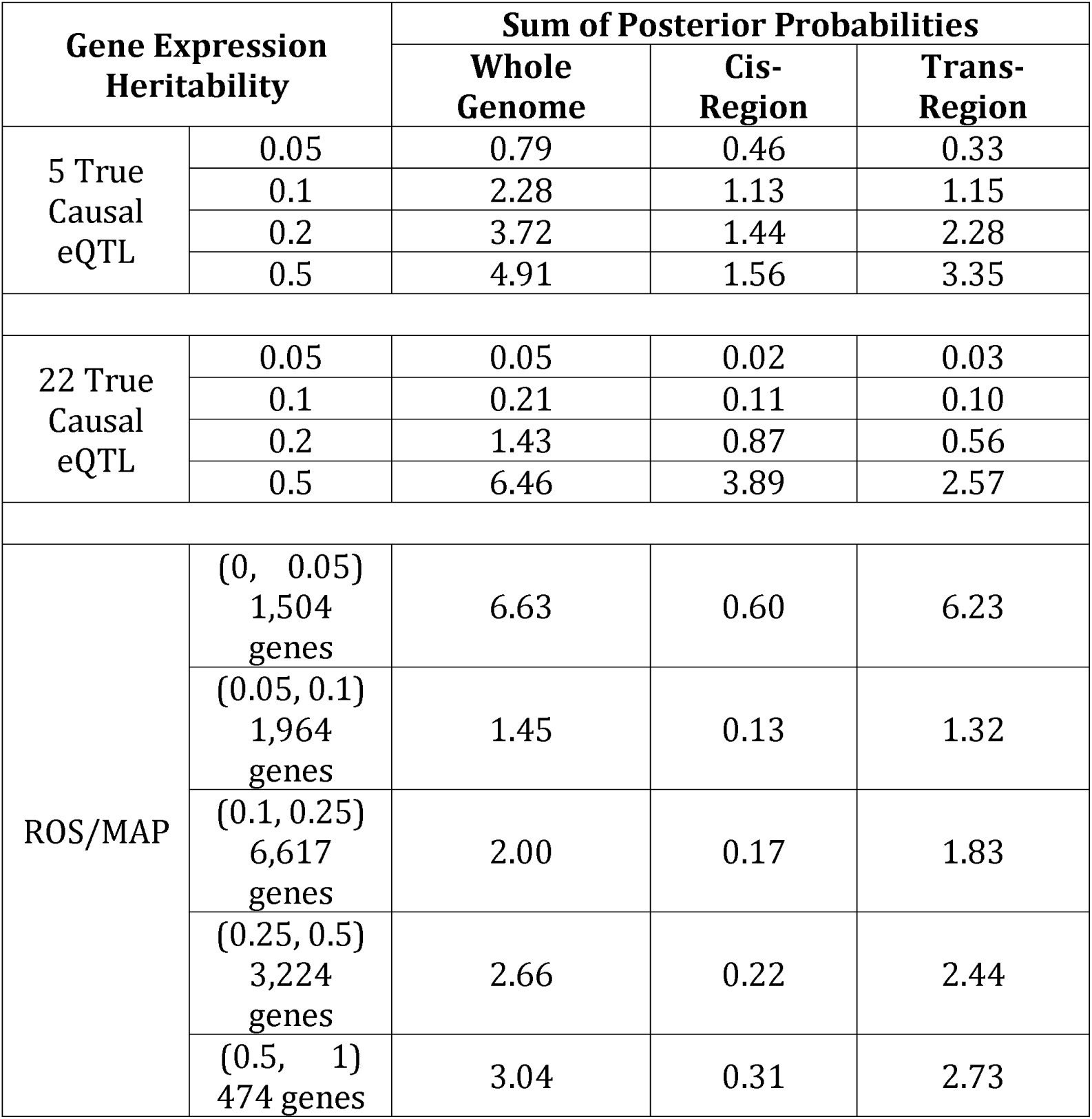
Average sums of posterior probabilities of having non-zero eQTL effect sizes that are stratified based on gene expression heritability (either true simulated heritability in simulation studies or the range of train R^2^ of the fitted BVSR models with ROS/MAP data). The simulation scenarios presented here are those with 2 of 5 and 11 of 22 true causal eQTL from the cis- regions.

| Gene Expression Heritability |  | Sum of Posterior Probabilities |  |  |
| --- | --- | --- | --- | --- |
|  |  | Whole Genome | Cis-Region | Trans-Region |
| 5 True Causal eQTL | 0.05 | 0.79 | 0.46 | 0.33 |
|  | 0.1 | 2.28 | 1.13 | 1.15 |
|  | 0.2 | 3.72 | 1.44 | 2.28 |
|  | 0.5 | 4.91 | 1.56 | 3.35 |
| 22 True Causal eQTL | 0.05 | 0.05 | 0.02 | 0.03 |
|  | 0.1 | 0.21 | 0.11 | 0.10 |
|  | 0.2 | 1.43 | 0.87 | 0.56 |
|  | 0.5 | 6.46 | 3.89 | 2.57 |
| ROS/MAP | (0, 0.05)<br>1,504 genes | 6.63 | 0.60 | 6.23 |
|  | (0.05, 0.1)<br>1,964 genes | 1.45 | 0.13 | 1.32 |
|  | (0.1, 0.25)<br>6,617 genes | 2.00 | 0.17 | 1.83 |
|  | (0.25, 0.5)<br>3,224 genes | 2.66 | 0.22 | 2.44 |
|  | (0.5, 1)<br>474 genes | 3.04 | 0.31 | 2.73 |

For 14,156 genes with fitted GReX prediction models by BVSR using the ROS/MAP data, after excluding 19 outlier genes with >100 expected eQTL, we obtained the average number of expected eQTL as 2.44 (SD = 5.70) across genome-wide regions, 0.25 (SD = 1.24) for cis-eQTL, and 2.48 (SD = 5.49) for trans-eQTL. That is, on overage, 88% of eQTL were from trans- genome regions with respect to the target gene. We can see that ∼90% genes with train R^2^ > 0.05 have ∼2-3 average expected eQTL, and ∼10% genes with train R^2^ < 0.05 have >5 average expected eQTL (Table 4). By linking these findings with our simulation studies where train ^2^ is likely to be >0.05 when true expression heritability is >0.1, we can conclude that ∼90% genes are likely to have true expression heritability >0.1.

Additionally, from our Bayesian estimates of the cis- and trans- specific posterior probabilities of having non-zero eQTL effect sizes (i.e., *π*_*cis*_, *π*_*trans*_ in Equation 3) for genome-wide genes using ROS/MAP data (Figure S11), we can see that *π*_*cis*_ and *π*_*trans*_ clearly follow different distributions. This also validates our assumptions of respective prior distribution for cis- and trans- hyper parameters.

## Discussion

In this paper, we proposed and validated a Bayesian Genome-wide TWAS (BGW-TWAS) method based on the BVSR^24^ model to leverage the information of both cis- and trans- eQTL. We derived an efficient computational approach to fit the BVSR model with large-scale genomic data, by pruning genome regions that contain either at least one cis-SNP or one potential trans-eQTL and adapting the previously developed scalable EM-MCMC algorithm^25^ with pre-calculated LD coefficients and summary statistics from standard eQTL analyses. BGW-TWAS extends previous TWAS methods^11-13^ that only utilize partial genotype information from a small window of cis*-* SNPs to train the GReX imputation model.

Genotype data of trans-eQTL have been shown to explain a significant amount of variation of expression quantitative traits and provide important molecular mechanisms underlying known GWAS loci of complex diseases^22; 23^. The results from our simulation and application studies demonstrated that BGW-TWAS improves the yield of TWAS by levering both cis- and trans- eQTL information. For example, higher precision of GReX prediction and power of TWAS were obtained in our simulation studies when true causal trans-eQTL existed. These results showed that BGW-TWAS has a greater advantage for scenarios where eQTL have relatively large effect sizes for the expression quantitative traits (e.g., 5 vs. 22 true causal eQTL with the same expression heritability). This is because variable selection by the BVSR model is designed to select sparse signals with relatively large effect sizes as shown in previous GWAS^24; 25^.

By applying our Bayesian approach to several human AD datasets, we identified a risk gene (*ZC3H12B*) with GReX values that were significantly associated with both clinical diagnosis of AD and postmortem AD pathology indices (neurofibrillary tangle density and global measure of AD pathology). This association was not identified by existing TWAS methods because this gene is shown to be completely driven by trans-eQTL. Importantly, a potential biological mechanism was revealed by showing that the top driven trans-eQTL of *ZC3H12B* are known GWAS signals of AD^43^ and blood lipids^44-46^ and <12KB away from the famous well-known AD risk gene (*APOE*)^47^. Thus, we expect BGW-TWAS leveraging both cis- and trans-eQTL has potential for making a large impact on advancing our understanding of complex human diseases and traits.

By fitting BVSR models using both cis- and trans- eQTL, we can not only account for the uncertainty for a SNP to be an eQTL to predict GReX (Equation 5), but also use the sum of posterior probabilities of having non-zero eQTL effect sizes to estimate the expected number of eQTL^24; 25^. The distribution of expected eQTL can also help characterize the underlying genetic architecture of expression quantitative traits.

The current study has several limitations. First, while BGW-TWAS reduces the computational burden for modeling both cis- and trans- eQTL, its computing costs are still substantial to train GReX prediction models for genome-wide genes (∼20K) per tissue type. It requires approximately 30 minutes of computation time and 3GB memory per gene (with parallel computation implemented in 4 CPU cores). Parallel computation can be employed to make use of high-performance computation clusters with multiple cores to reduce computation time. Second, our current method is designed to use pre-calculated in-sample LD coefficients and summary statistics from single variant eQTL analyses, further work is required to expand this approach to use approximate LD coefficients generated from reference samples of the same ethnicity. Third, our simulation studies showed that the non-parametric Bayesian method TIGAR performed best when all causal eQTL are cis- with relatively small effect sizes (e.g., 22 true causal cis-eQTL). Our TWAS results of AD using the IGAP summary statistics demonstrated that TIGAR and BGW-TWAS yield complementary findings. These results highlight the potential utility of leveraging both methods especially for studies in which the true distributions of cis- and trans- eQTL of the test genes are generally unknown.

In conclusion, the BGW-TWAS method presented herein provides a framework for leveraging information from both cis- and trans- eQTL to conduct gene-based association studies. Because trans- QTL are common for other quantitative omics traits, e.g., epigenetic, proteomic, and metabonomic, our proposed computational procedure would be to investigate other quantitative omics traits in gene-based association studies. Integrative method developments will stand to benefit from our BGW-TWAS method, especially the perspectives of leveraging information from trans- QTL and efficient computation techniques derived from this paper. In addition, BGW-TWAS can be applied to study other complex human phenotypes to identify potential risk genes that could be targeted in further drug discovery.

## Supplemental Data

Supplemental Figures, Tables, Methods, and References.

## Data and Code Availability

ROS/MAP data can be requested through Rush Alzheimer’s Disease Center (http://www.radc.rush.edu/) and synapse (https://www.synapse.org/#!Synapse:syn3219045). MCADGS data can be requested through synapse (https://www.synapse.org/#!Synapse:syn2910256). IGAP summary statistics are available from http://web.pasteur-lille.fr/en/recherche/u744/igap/igap_download.php. Summary statistics generated from our BGW-TWAS methods for studying AD are publicly available through synapse (https://www.synapse.org/BGW_TWAS). Source code of BGW-TWAS is available through Github with link https://github.com/yanglab-emory/BGW-TWAS.

## Acknowledgement

J.Y. was supported by the startup funding from Department of Human Genetics at Emory University School of Medicine. ROS/MAP study data were provided by the Rush Alzheimer’s Disease Center, Rush University Medical Center, Chicago, IL. Data collection was supported through funding by NIA grants P30AG10161, R01AG15819, R01AG17917, R01AG30146, R01AG36836, U01AG32984, U01AG46152, U01AG61356, the Illinois Department of Public Health, and the Translational Genomics Research Institute. The MCADGC led by Dr. Nilüfer Ertekin-Taner and Dr. Steven G. Younkin, Mayo Clinic, Jacksonville, FL uses samples from the Mayo Clinic Study of Aging, the Mayo Clinic Alzheimer’s Disease Research Center, and the Mayo Clinic Brain Bank. MCADGC data collection was supported through funding by NIA grants P50 AG016574, R01 AG032990, U01 AG046139, R01 AG018023, U01 AG006576, U01 AG006786, R01 AG025711, R01 AG017216, R01 AG003949, NINDS grant R01 NS080820, CurePSP Foundation, and support from Mayo Foundation.

## Web Resources

BGW-TWAS, https://github.com/yanglab-emory/BGW-TWAS

TIGAR, https://github.com/yanglab-emory/TIGAR

PrediXcan, https://github.com/hakyim/PrediXcan

