## Supplemental Note for "Bayesian Genome-wide TWAS method to leverage both cis- and trans- eQTL information through summary statistics"

### 1 Supplemental Figures

#### 1.1 Training $R^2$ comparison

**Figure S** 1: Compare train $R^2$ obtained by our BGW method, PrediXcan, and TIGAR.

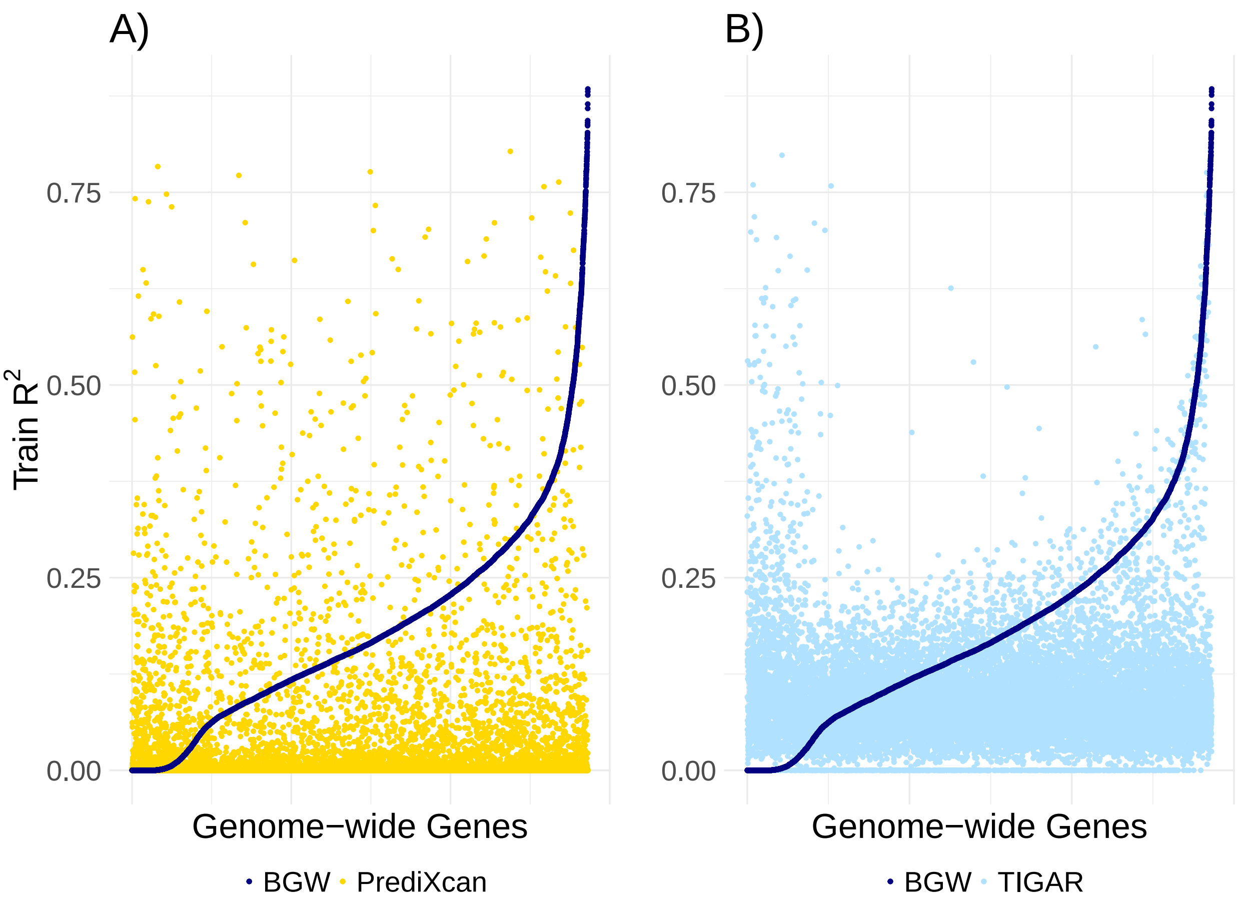


Train $R^2$ value per gene obtained by our BGW method (dark blue), PrediXcan (yellow), and TIGAR (baby blue) for genome-wide genes are plotted, where genes were ranked in the increasing order of $R^2$ by BGW.

#### 1.2 TWAS results using individual-level GWAS data of ROS/MAP and MCADGS

**Figure S** 2: Manhattan plots of TWAS results of AD clinical diagnosis and global AD pathology by using BVSR cis-eQTL estimates only.

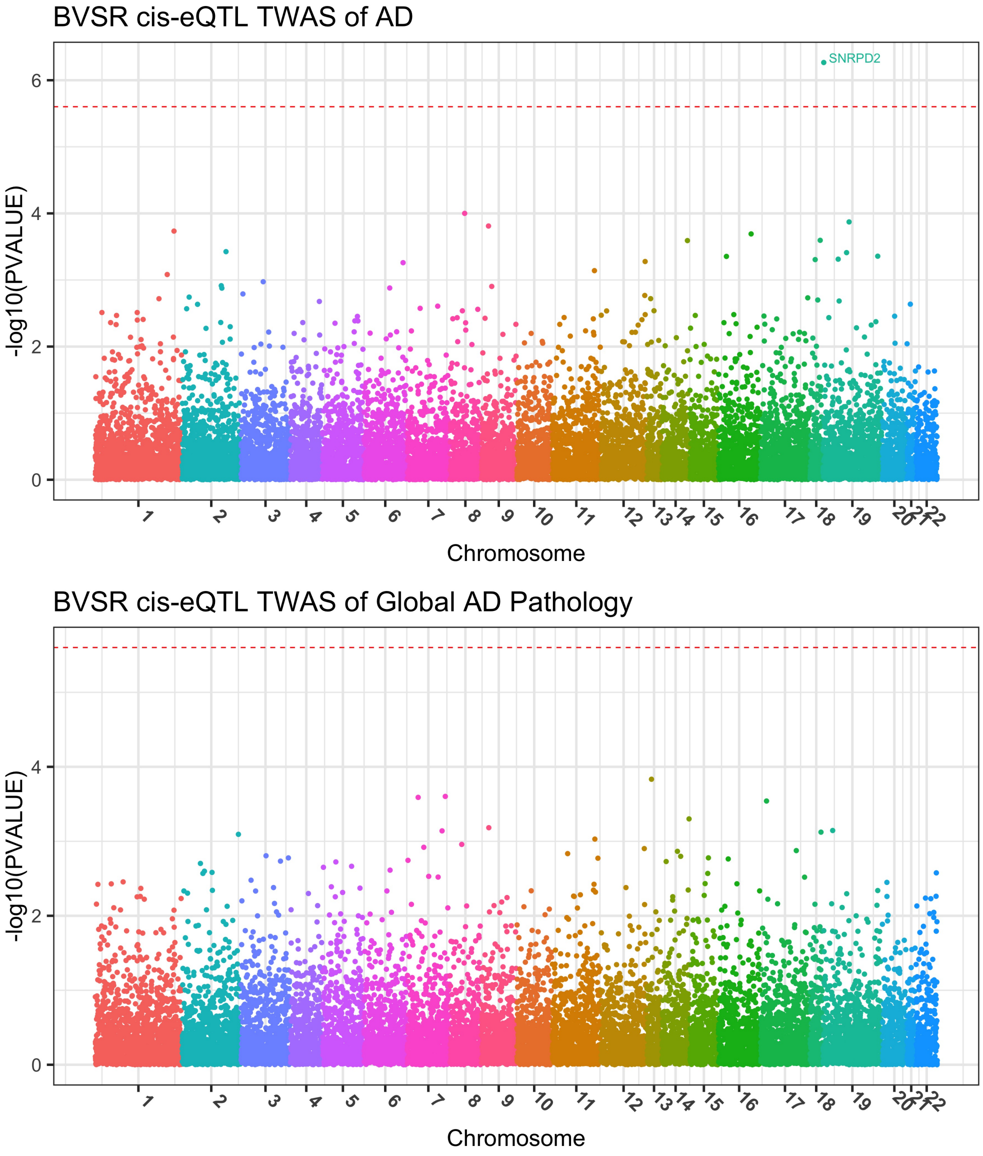

**Figure S** 3: Manhattan plots of TWAS results of neurofibrillary tangle density (tangles) and $\beta$-amyloid load (amyloid) by using BVSR cis-eQTL estimates only.

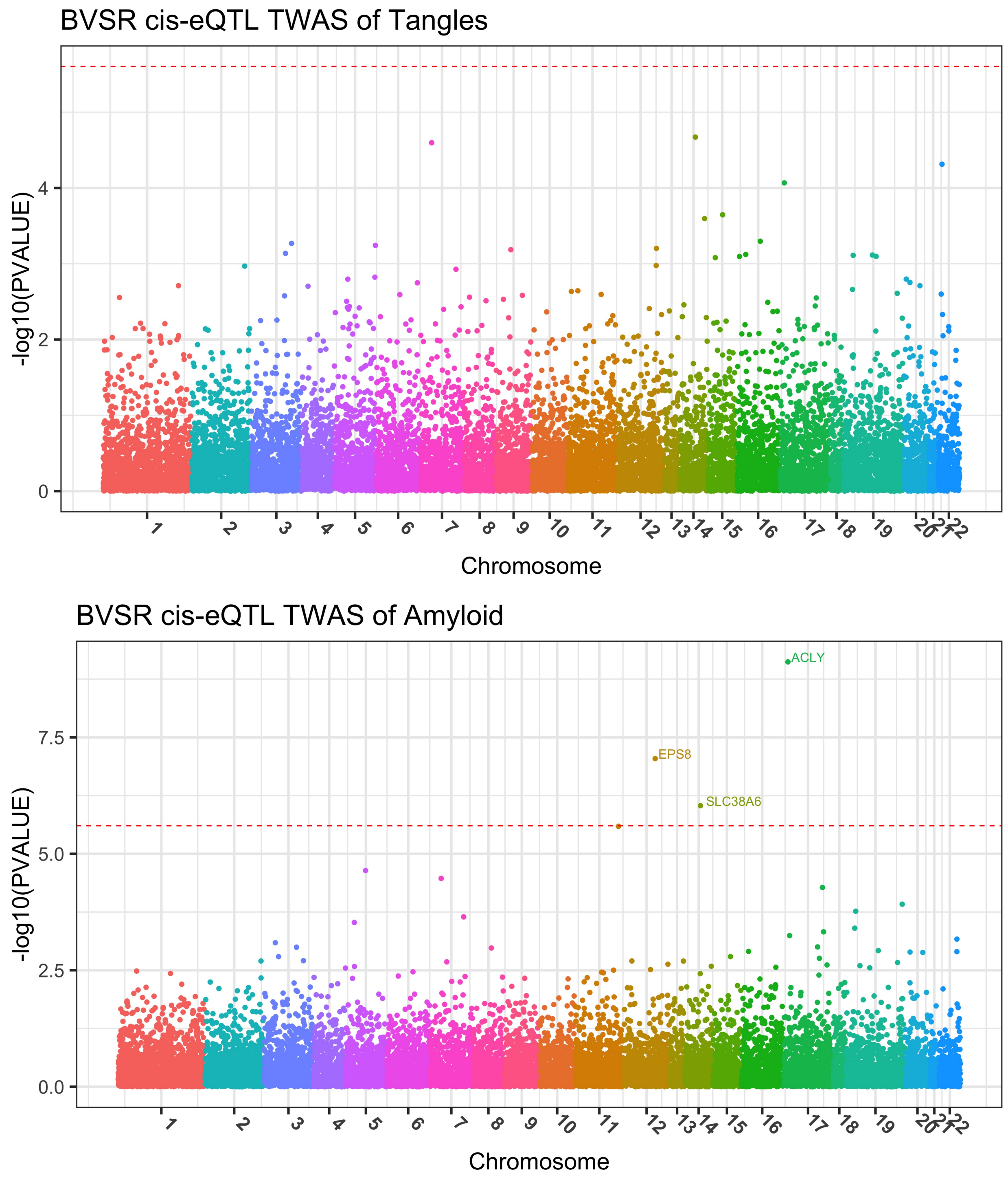

**Figure S** 4: Manhattan plots of TWAS results of AD clinical diagnosis by PrediXcan and TIGAR.

**TWAS of AD by PrediXcan**

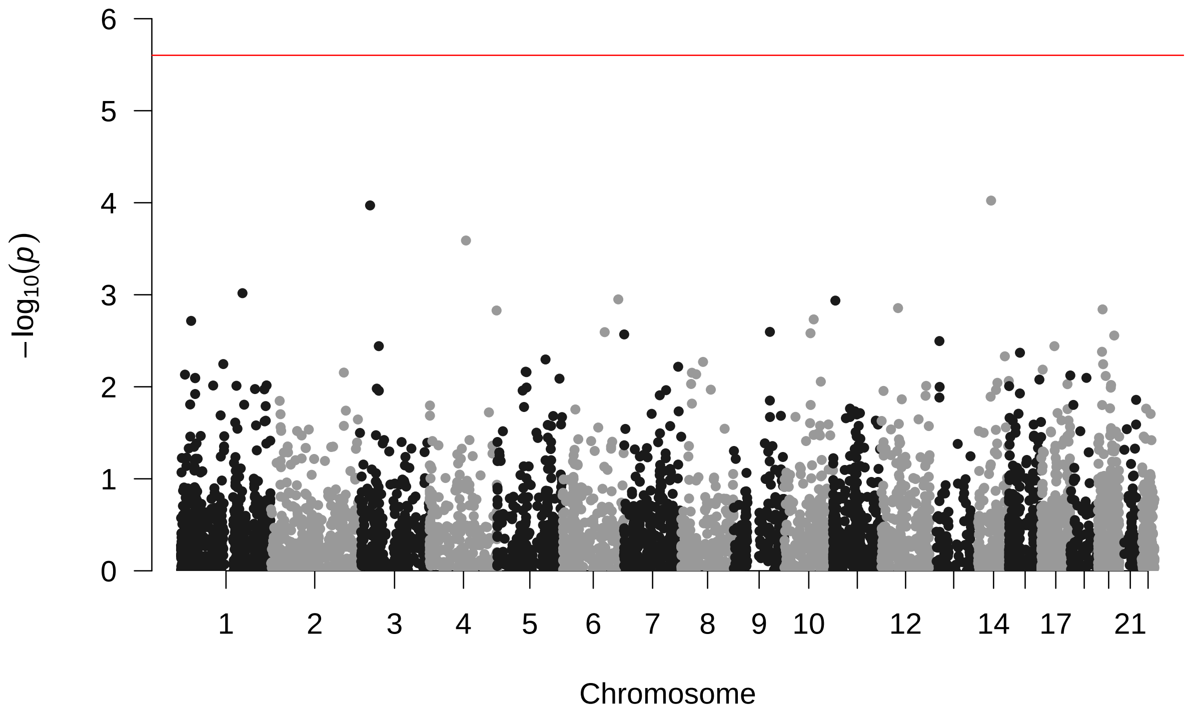


**TWAS of AD by TIGAR**

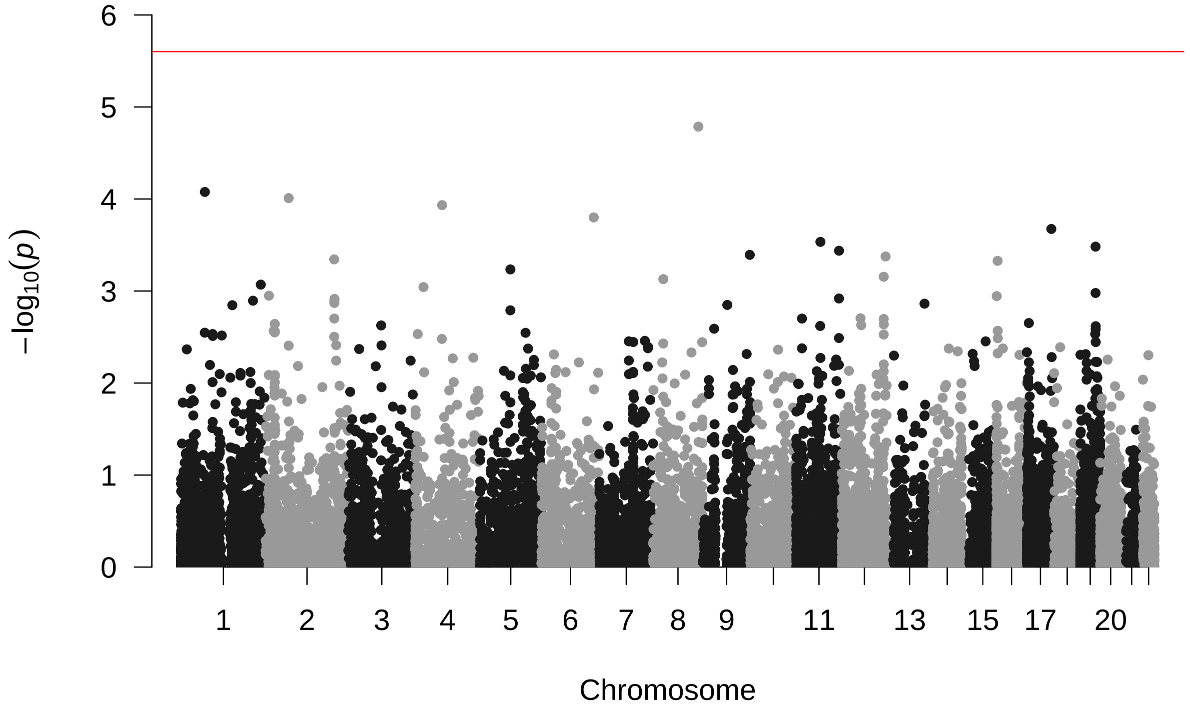

**Figure S** 5: Manhattan plots of TWAS results of global AD pathology by PrediXcan and TIGAR.

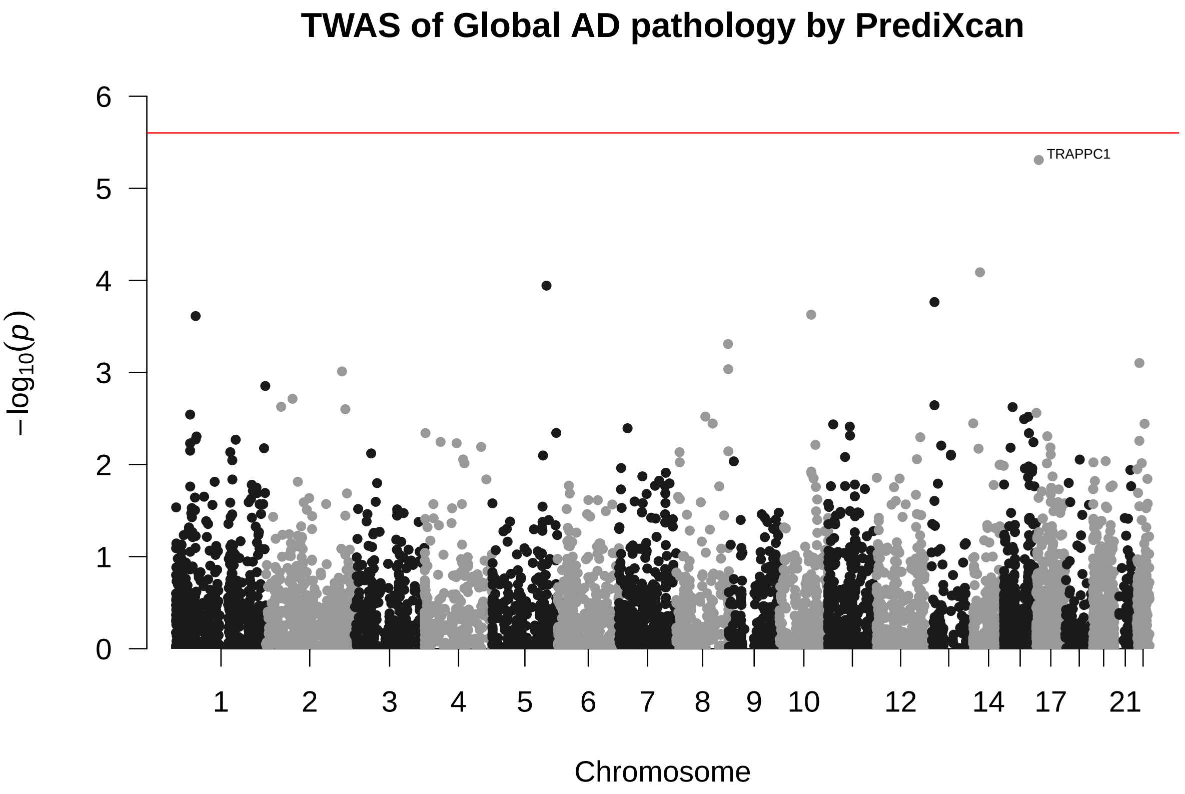


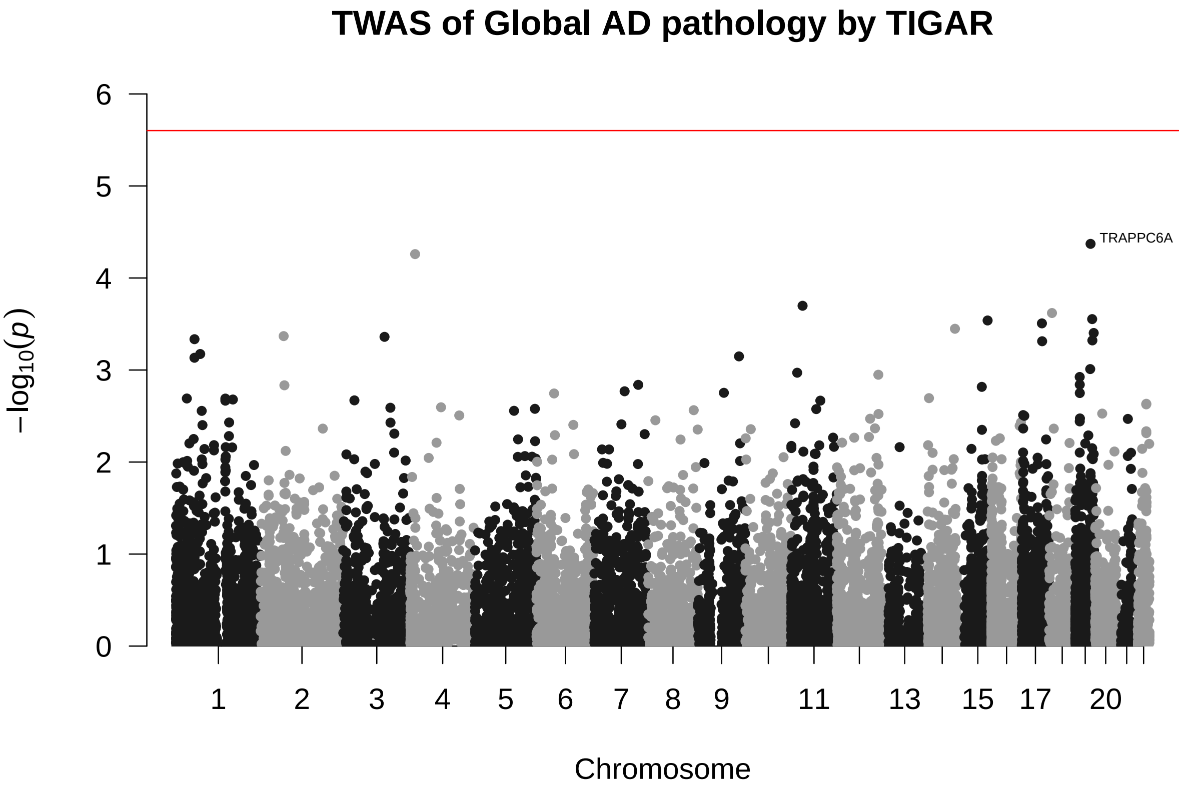

**Figure S** 6: Manhattan plots of TWAS results of neurofibrillary tangle density (tangles) by PrediXcan and TIGAR.

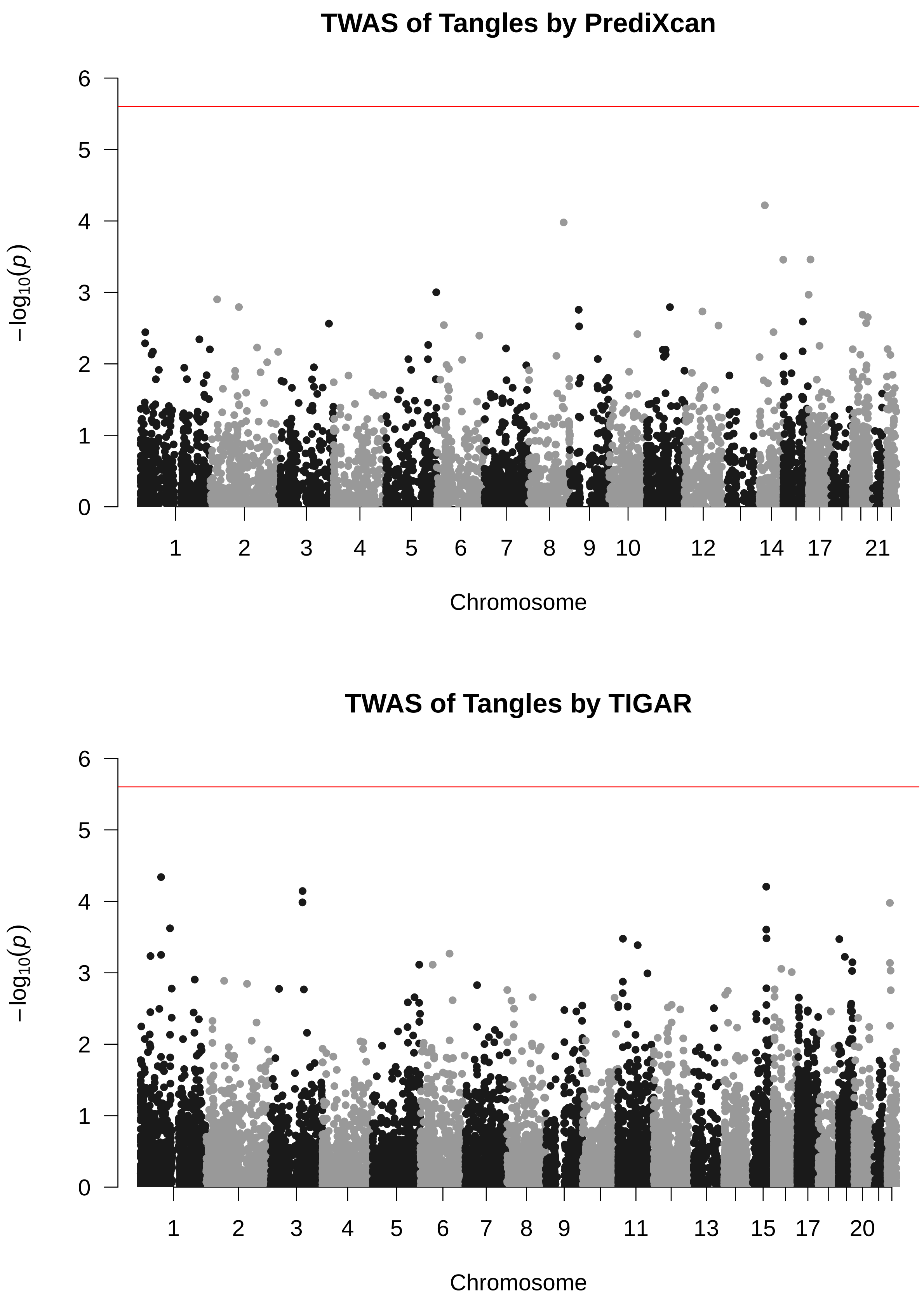

**Figure S 7**: Manhattan plots of TWAS results of $\beta$-amyloid load (amyloid) by PrediXcan and TIGAR.

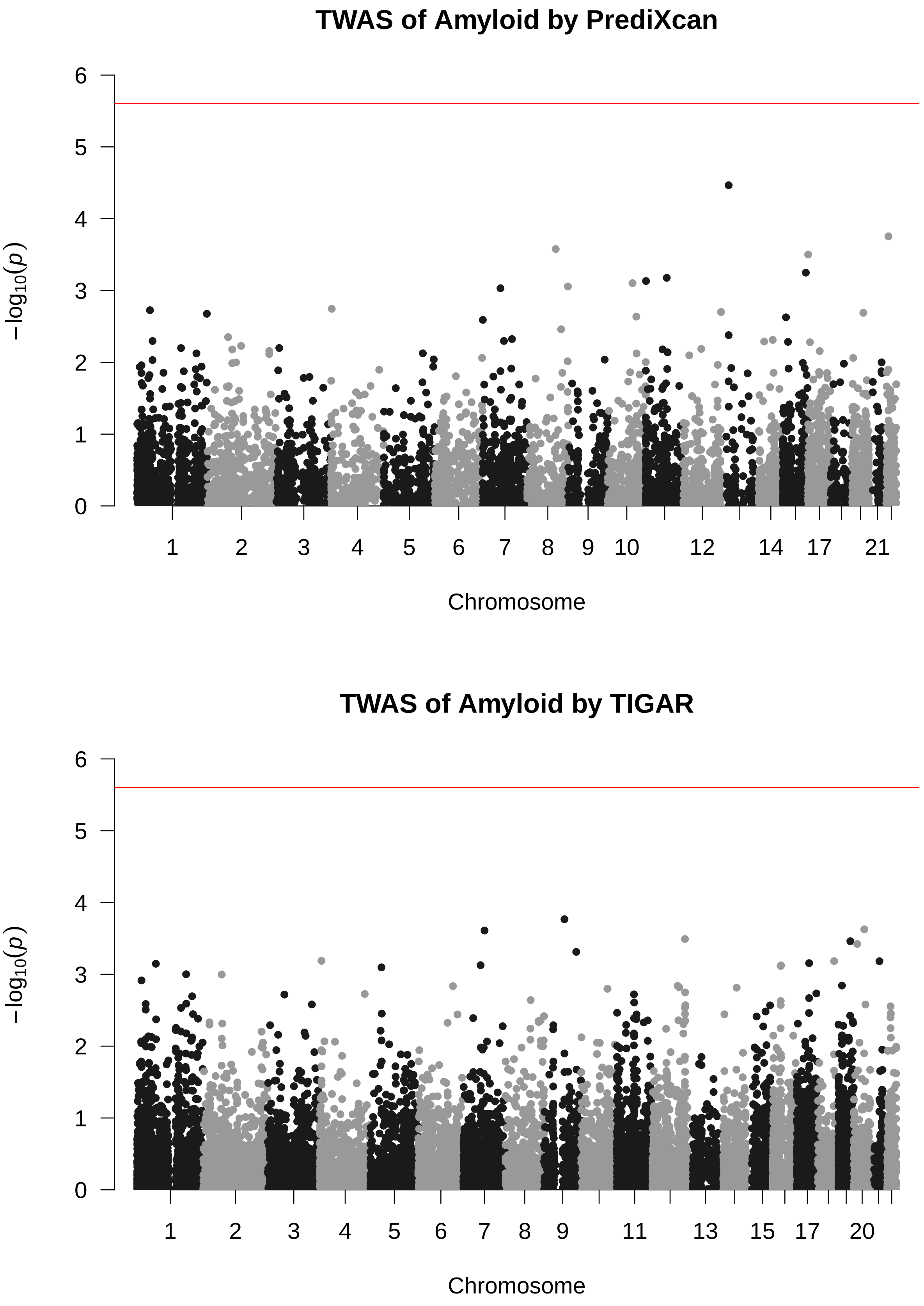

**Figure S** 8: Manhattan plots of standard eQTL analysis results of genes *ZC3H12B* and *KCTD12* by single variant tests.

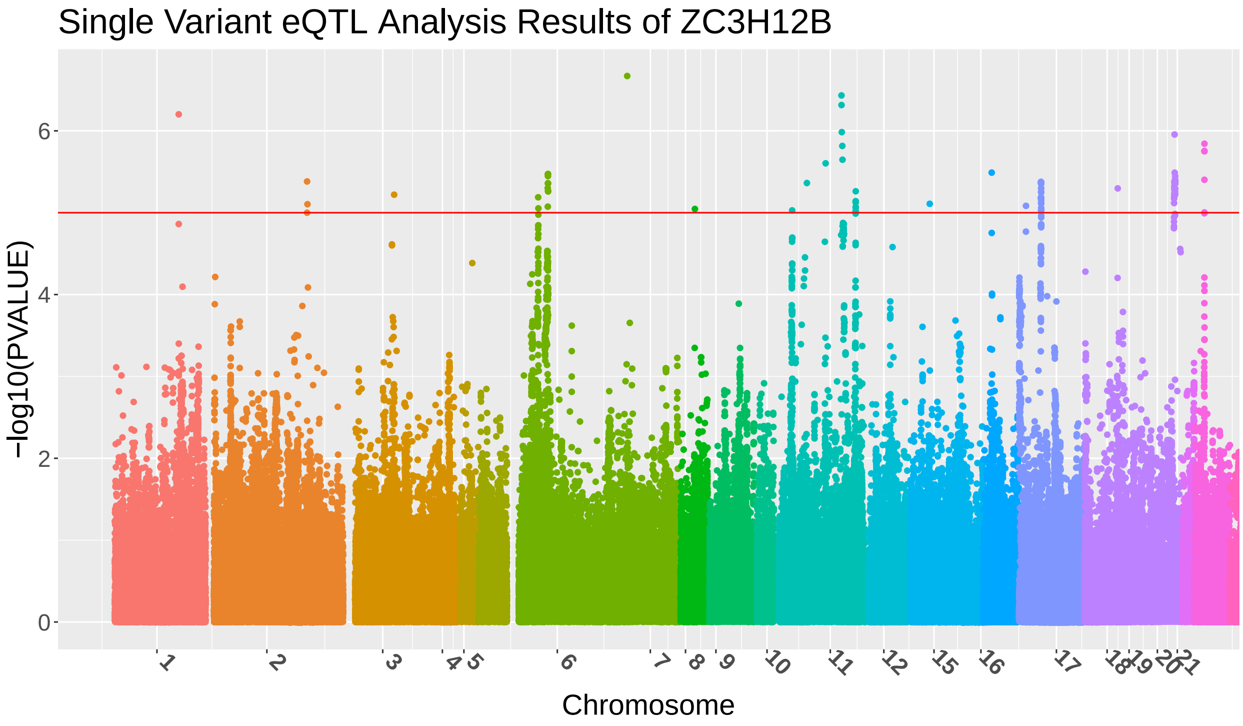


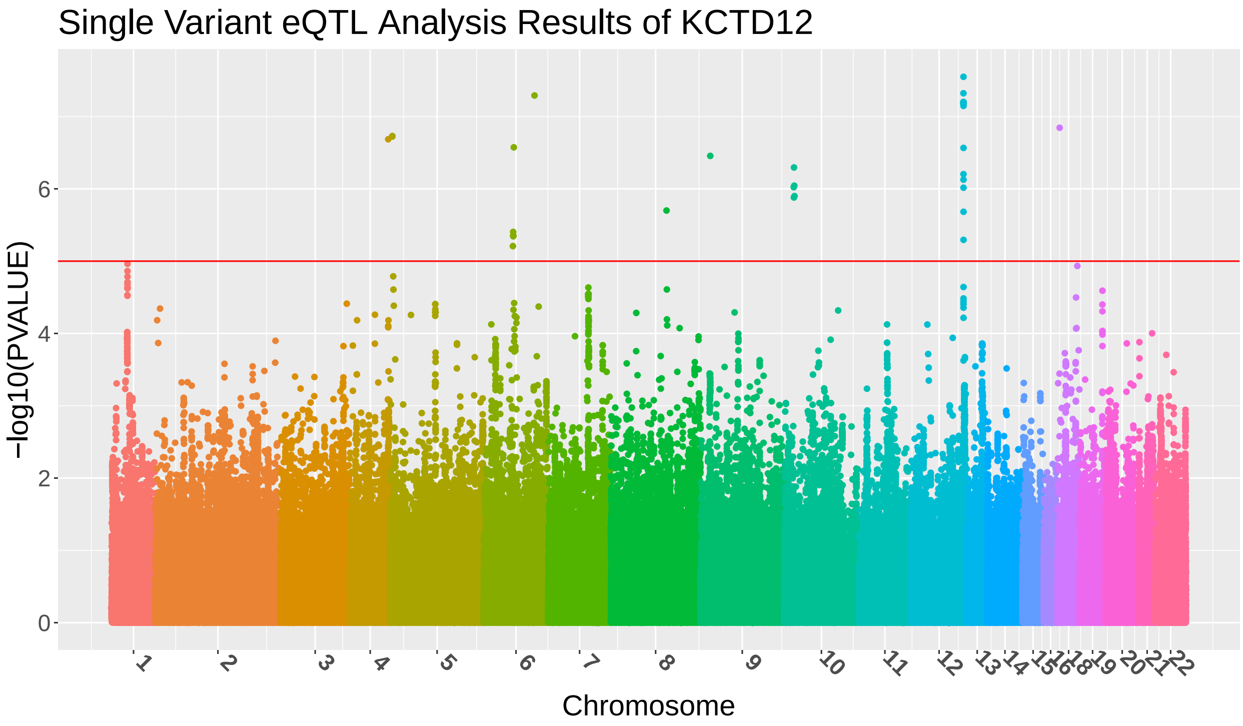

#### 1.3 TWAS results using IGAP summary-level GWAS data of AD

**Figure S** 9: Manhattan plots of TWAS results using IGAP summary statistics of AD by using both cis-eQTL and trans-eQTL (BGW-TWAS) and only cis-eQTL estimates obtained by BVSR.

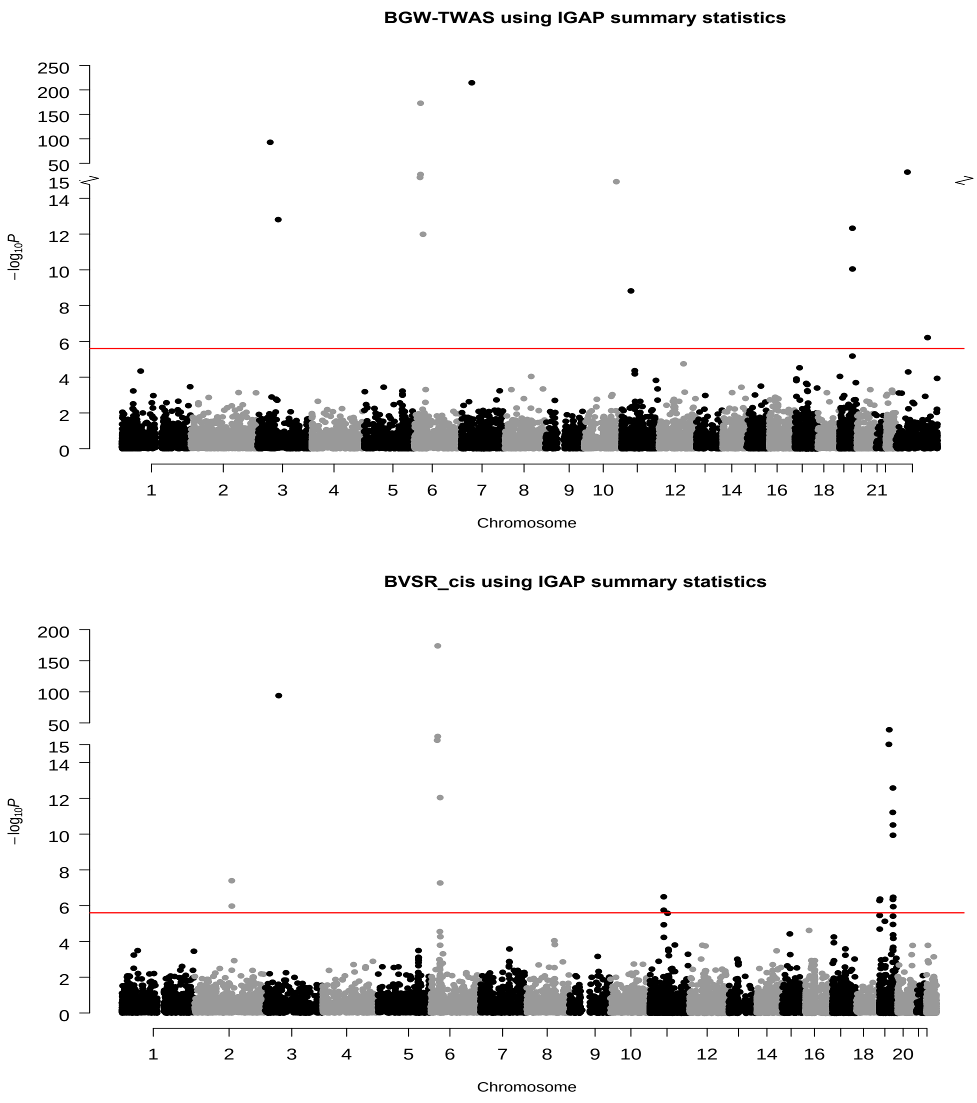

**Figure S** 10: Manhattan plots of TWAS results using IGAP summary statistics of AD by S-PrediXcan and TIGAR .

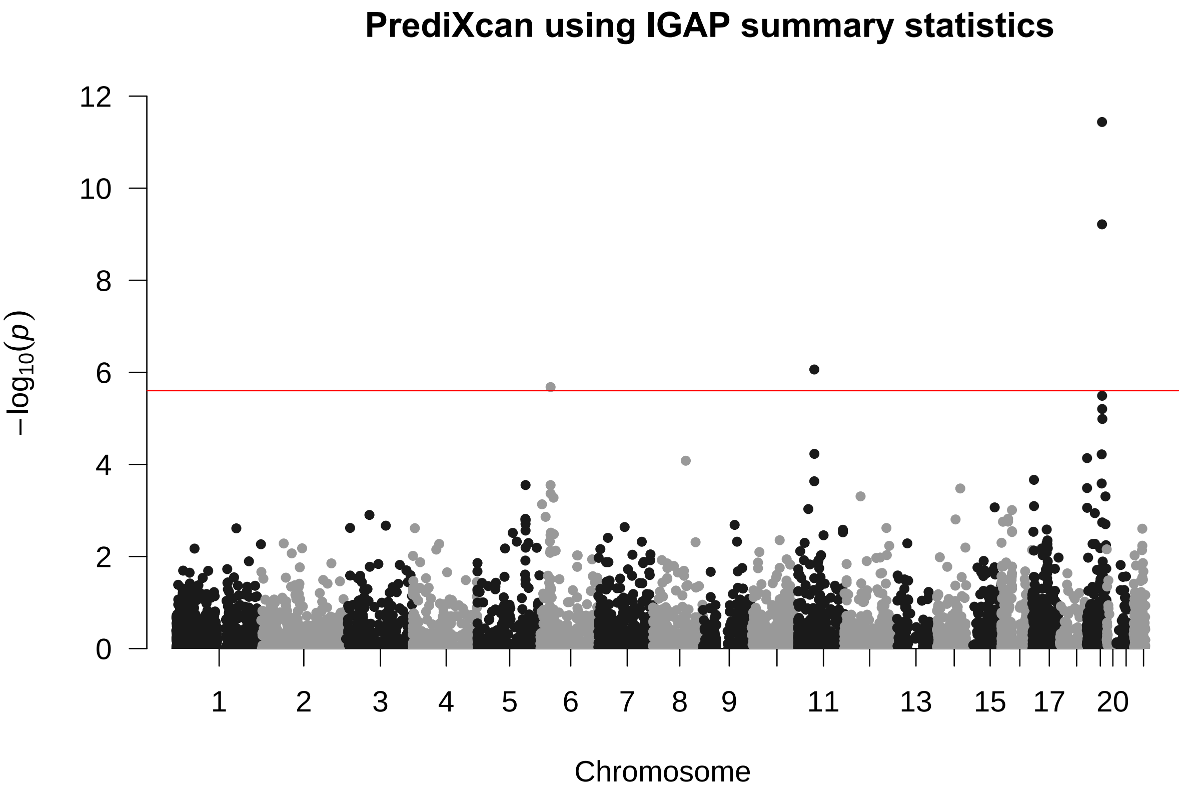


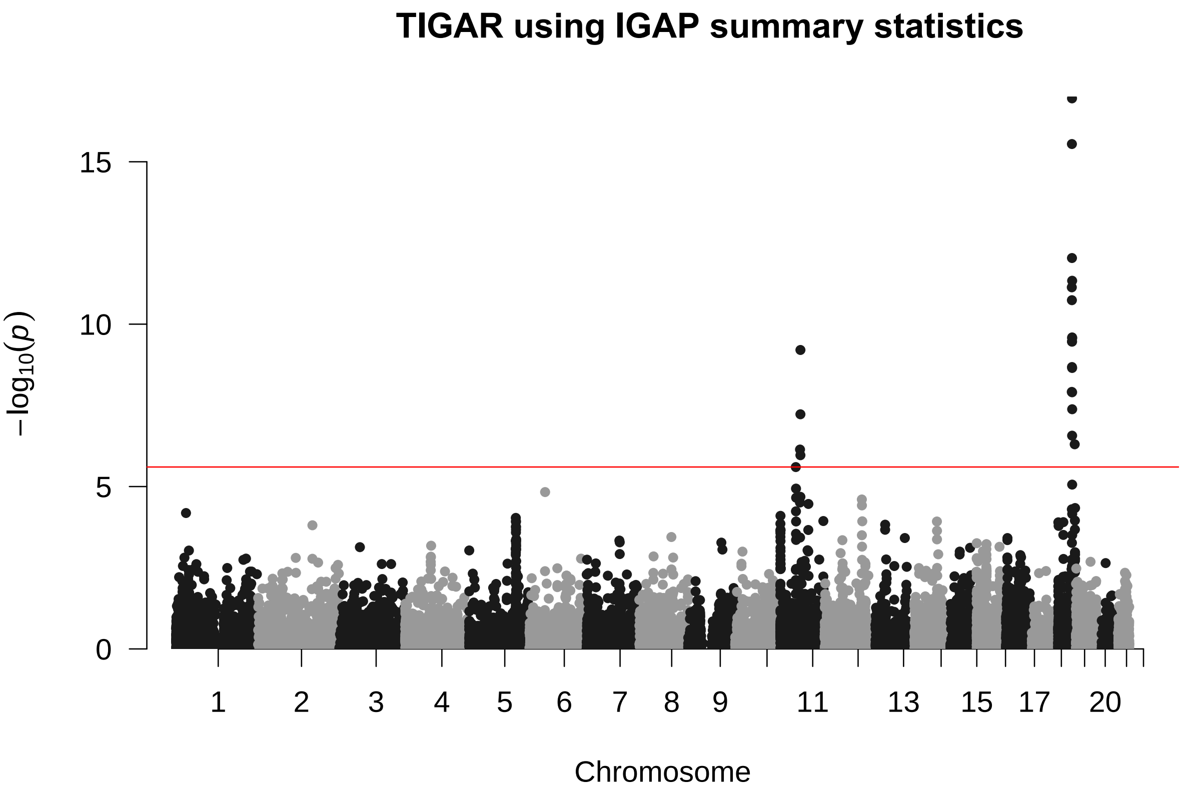

**Figure S** 11: Histogram of $log_{10}$ Bayesian estimates for cis- and trans- specific probabilities of being an eQTL ($\pi_{cis}, \pi_{trans}$), over genome-wide genes.

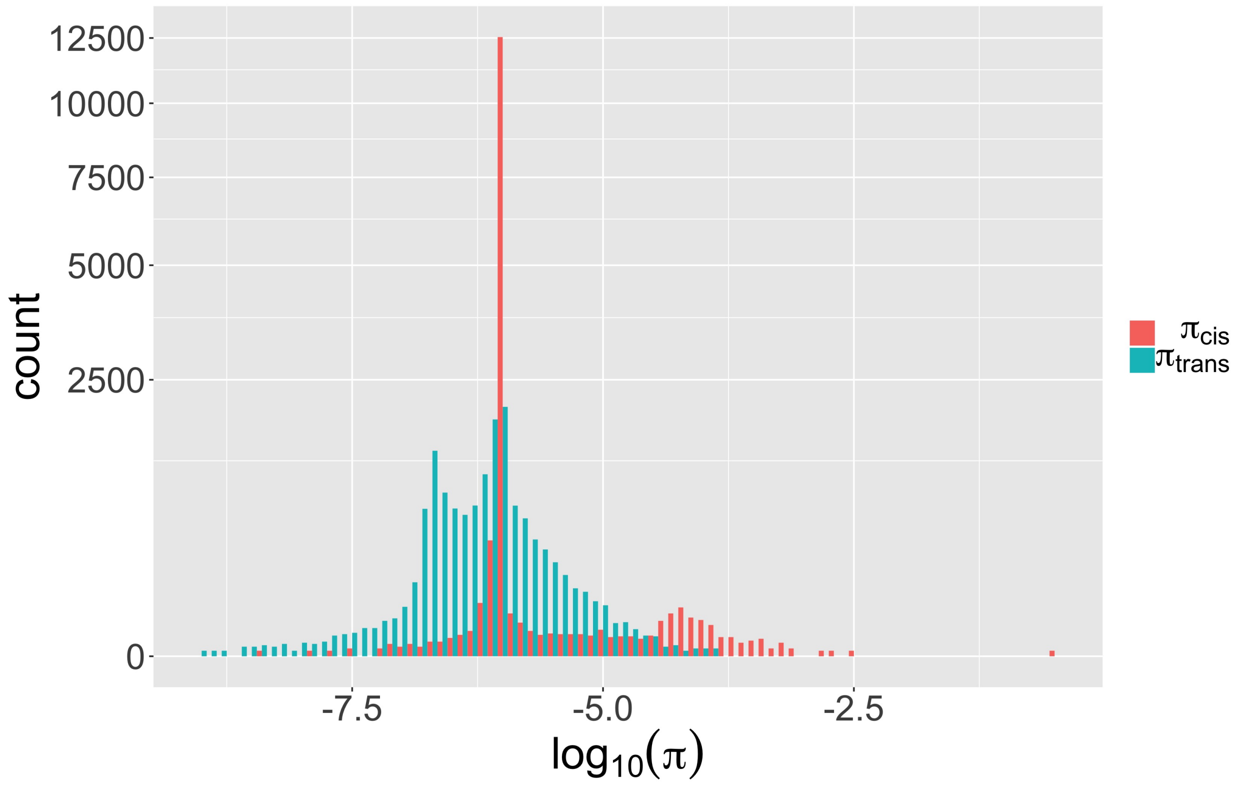

#### 2 Supplemental Tables

| Gene | CHR | Position | P-VALUE | Z-score |
|---|---|---|---|---|
| *WDR33* | 2 | 128,458,595 | 4.01e-08 | -5.49 |
| *SAP130* | 2 | 128,698,790 | 1.05e-06 | -4.88 |
| *GPX1*[a] | 3 | 493,96,033 | 2.45e-98 | 21.04 |
| *BTN3A2*[a] | 6 | 26,365,386 | 1.56e-26 | 10.66 |
| *ZNF192*[a] | 6 | 28,109,715 | 1.25e-32 | -11.89 |
| *AL022393.7*[a] | 6 | 28,143,965 | 2.24e-178 | -28.47 |
| *HLA-DRB1*[a,b] | 6 | 32,546,545 | 8.99e-13 | 7.14 |
| *HLA-DQB1* | 6 | 32,627,243 | 5.33e-08 | -5.43 |
| *RP11-750H9.5* | 11 | 4,740,4698 | 3.08e-07 | -5.11 |
| *SLC39A13*[b] | 11 | 47,428,682 | 1.73e-06 | -4.78 |
| *FSTL3* | 19 | 676,388 | 4.98e-07 | -5.02 |
| *MKNK2* | 19 | 2,037,480 | 4.12e-07 | -5.06 |
| *ZNF227*[c] | 19 | 44,716,690 | 5.87e-12 | 6.88 |
| *ZFP112*[c] | 19 | 44,830,705 | 5.79e-20 | 9.14 |
| *PVR*[a,b,c] | 19 | 45,147,097 | 4.3e-07 | -5.05 |
| *CEACAM19*[a,b,c] | 19 | 45,174,723 | 2.54e-13 | 7.31 |
| *TOMM40* | 19 | 45,394,476 | 2.97e-11 | -6.64 |
| *APOC1*[a] | 19 | 45,417,920 | 1.11e-10 | 6.45 |
| *FOSB* | 19 | 45,971,252 | 3.26e-07 | -5.10 |
| *SNRPD2* | 19 | 46,190,712 | 1.97e-43 | -13.81 |
| *FBXO46* | 19 | 46,213,886 | 1.09e-06 | -4.87 |

**Table S** 1: Significant genes identified by TWAS using BVSR cis-eQTL estimates only.
a. Genes that were also identified by BGW-TWAS.
b. Genes that were also identified by PrediXcan.
c. Genes that were also identified by TIGAR.

| Gene | CHR | Position | P-VALUE | Z-score |
|---|---|---|---|---|
| *HLA-DRB1*[a] | 6 | 32546545 | 2.06e-06 | 4.75 |
| *SLC39A13*[a] | 11 | 47428682 | 8.57e-07 | -4.92 |
| *PVR*[a,b] | 19 | 45147097 | 6.02e-10 | -6.19 |
| *CEACAM19*[a,b] | 19 | 45174723 | 3.6e-12 | 6.95 |

**Table S** 2: Significant genes identified by S-PrediXcan.

a. Genes that were also identified by BVSR estimates of cis-eQTL and trans-eQTL (BGW-TWAS) or cis-eQTL only.

b. Genes that were also identified by TIGAR.

| Gene | CHR | Position | P-VALUE | Z-score |
|---|---|---|---|---|
| *STX3* | 11 | 59,480,928 | 7.28e-07 | -4.95 |
| *PRPF19* | 11 | 60,658,201 | 5.98e-08 | 5.42 |
| *TMEM109* | 11 | 60,681,345 | 1.09e-06 | -4.88 |
| *TMEM132A* | 11 | 60,691,934 | 6.26e-10 | -6.18 |
| *ZNF227*[a] | 19 | 44,716,690 | 1.21e-08 | 5.7 |
| *ZFP112*[a] | 19 | 44,830,705 | 7.32e-12 | 6.85 |
| *PVR*[a,b] | 19 | 45,147,097 | 1.82e-11 | -6.72 |
| *CEACAM19*[a,b] | 19 | 45,174,723 | 2.83e-16 | 8.18 |
| *CLPTM1* | 19 | 45,457,847 | 9.17e-13 | -7.14 |
| *CLASRP* | 19 | 45,542,297 | 2.08e-09 | 5.99 |
| *ZNF296* | 19 | 45,574,758 | 3.45e-10 | -6.28 |
| *TRAPPC6A* | 19 | 45,666,186 | 1.11e-17 | -8.56 |
| *MARK4* | 19 | 45,754,549 | 3.41e-10 | 6.28 |
| *RTN2* | 19 | 45,988,549 | 2.7e-07 | 5.14 |
| *PPM1N* | 19 | 45,992,034 | 1.24e-08 | 5.69 |
| *OPA3* | 19 | 46,030,684 | 2.55e-10 | -6.32 |
| *EML2* | 19 | 46,112,659 | 2.22e-09 | 5.98 |
| *GIPR* | 19 | 46,171,501 | 4.58e-12 | 6.92 |
| *SNRPD2* | 19 | 46,190,712 | 2.69e-10 | 6.32 |
| *DMWD* | 19 | 46,286,204 | 4.15e-08 | -5.48 |
| *TTYH1* | 19 | 54,926,372 | 4.98e-07 | -5.03 |

**Table S** 3: Significant genes identified by TIGAR.
a. Genes that were also identified by BVSR estimates of cis-eQTL and trans-eQTL (BGW-TWAS) or cis-eQTL only.
b. Genes that were also identified by S-PrediXcan.

### 3 Supplemental Methods

#### 3.1 Bayesian Variable Selection Regression Model

Consider the following Bayesian variable selection regression (BVSR) model [1] for quantitative gene expression traits:

$$\mathcal{E}_{n\times 1} = \boldsymbol{X}_{n\times p}\boldsymbol{w}_{p\times 1} + \boldsymbol{\epsilon}_{n\times 1},\ w_i \sim \pi N(0, \sigma_w^2\sigma_\epsilon^2) + (1-\pi)\delta_0(\cdot),\ \epsilon_i \sim N(0,\sigma_\epsilon^2), \tag{1}$$

where $\mathcal{E}_{n\times 1}$ denotes the vector of centered quantitative expression levels for $n$ samples; $\boldsymbol{X}_{n\times p}$ denotes the centered genotype matrix of $p$ genetic variants; $\epsilon_i$ denotes the residual error independently and identically distributed (i.i.d.) with normal distribution $N(0,\sigma_\epsilon^2)$; and the broad sense "eQTL" effect size $w_i$ follows a spike-and-slab prior distribution [1, 2, 3] — that is, $w_i$ follows the normal distribution $N(0,\sigma_w^2\sigma_\epsilon^2)$ with probability $\pi$ and the point-mass density function $\delta_0(\cdot)$ at $0$ with probability $(1-\pi)$. Basically, $\delta_0(w_i) = 1$ if $w_i = 0$, otherwise $\delta_0(w_i) = 0$. The effect size variance $(\sigma_w^2\sigma_\epsilon^2)$ is assumed to be scaled by the error term variance $(\sigma_\epsilon^2)$ for the purpose of computational convenience.

As is typical of SNP-based genome-wide analyses, the genotype matrix $\boldsymbol{X}_{n\times p}$ contains either dosage data within range $[0,2]$ or genotype data with values $\{0,1,2\}$ denoting the expected or genotyped number of minor alleles. The assumption of the spike-and-slab prior for $w_i$ enforces variable selection in the regression model (1). Assuming both $\mathcal{E}_{n\times 1}$ and columns of $\boldsymbol{X}_{n\times p}$ are centered, the intercept term is omitted from the regression model.

##### 3.1.1 Model cis- and trans- eQTL

In this paper, we employ this BVSR model (1) to account for both cis- and trans- eQTL genotype data (cis- and trans- are defined based on SNP proximity to the target gene) for modeling quantitative gene expression traits. Particularly, we extend the BVSR model to allow for respective prior distributions for the effect sizes of cis- and trans- SNPs (i.e., eQTL) as follows:

$$\begin{aligned}
\mathcal{E}_g &= \boldsymbol{X}_{cis}\boldsymbol{w}_{cis} + \boldsymbol{X}_{trans}\boldsymbol{w}_{trans} + \boldsymbol{\epsilon} \\
w_{cis,i} &\sim \pi_{cis}N(0,\sigma_{cis}^2\sigma_\epsilon^2) + (1-\pi_{cis})\delta_0(w_{cis,i}) \\
w_{trans,i} &\sim \pi_{trans}N(0,\sigma_{trans}^2\sigma_\epsilon^2) + (1-\pi_{trans})\delta_0(w_{trans,i}), \\
\epsilon_i &\sim N(0,\sigma_\epsilon^2).
\end{aligned} \tag{2}$$

Generally, SNPs within $\pm$1MB of the flanking 5' and 3' ends of the target gene are considered as cis-SNPs, and other SNPs on the genome are considered as trans-SNPs.

Note that cis- and trans- can be viewed as two non-overlapped annotations for SNPs in the BVSR model (1), which makes model (2) a special case of the previously developed Bayesian Functional GWAS (BFGWAS) method [4]. Similarly, the following independent and conjugate hyper priors are assumed for hyper parameters in model (2):

$$\begin{aligned} \pi_{cis} &\sim Beta(a_{cis}, b_{cis}),\ \sigma^2_{cis} \sim IG(k_1, k_2), \\ \pi_{trans} &\sim Beta(a_{trans}, b_{trans}),\ \sigma^2_{trans} \sim IG(k_3, k_4), \\ \sigma^2_{\epsilon} &\sim IG(k_5, k_6) \end{aligned} \tag{3}$$

where $Beta(a_q, b_q)$ denotes a Beta distribution with positive shape parameters $a_q$ and $b_q$ for $q = \{cis, trans\}$, and an Inverse-Gamma distribution $IG(k_s, k_l)$ with shape parameters $k_s$ and scale parameters $k_l$ is assumed for $(\sigma^2_{cis}, \sigma^2_{trans}, \sigma^2_{\epsilon})$ with respective shape and scale parameters.

Hyper prior values are chosen for $a_q$ and $b_q$ to enforce a sparse model, such that the mean of the Beta distribution $\frac{a_q}{a_q+b_q} = 10^{-6}$ with $(a_q + b_q)$ equal to the total number of variants of respective annotation $q = \{cis, trans\}$. The hyper priors for Inverse Gamma are taken as $k_1 = k_2 = k_3 = k_4 = k_5 = k_6 = 0.1$ to induce non-informative priors for $(\sigma^2_{cis}, \sigma^2_{trans}, \sigma^2_{\epsilon})$. Thus, the posterior estimates of $(\pi_{cis}, \sigma^2_{cis}, \pi_{trans}, \sigma^2_{trans})$ will mainly be driven by data likelihood.

##### 3.1.2 Latent Indicator Variable

To facilitate computation, a latent indicator vector $\boldsymbol{\gamma}_{p\times 1}$ is introduced [2] into the model (2), where each element $\gamma_i \in \{0, 1\}$ indicates whether the corresponding $i$th effect $w_{q,i}$ equals to 0 with $\gamma_i = 0$ or follows the $N(0, \sigma^2_q \sigma^2_{\epsilon})$ distribution with $\gamma_i = 1$. Equivalently,

$$\gamma_i \sim Bernoulli(\pi_i),\ \boldsymbol{w}_{-\boldsymbol{\gamma}} \sim \delta_0(\cdot),\ \boldsymbol{w}_{\boldsymbol{\gamma}} \sim MVN_{|\boldsymbol{\gamma}|}(0, \sigma^2_{\epsilon}\boldsymbol{V}_{\boldsymbol{\gamma}}), \tag{4}$$

where $|\boldsymbol{\gamma}|$ denotes the number of non-zero entries in $\boldsymbol{\gamma}$; $\boldsymbol{w}_{-\boldsymbol{\gamma}}$ denotes the sub-vector of $\boldsymbol{w}_{p\times 1}$ corresponding to variants with $\gamma_i = 0$; $\boldsymbol{w}_{\boldsymbol{\gamma}}$ denotes the sub-vector of $\boldsymbol{w}_{p\times 1}$ corresponding to the variants with $\{\gamma_j = 1; j = 1, \cdots, |\boldsymbol{\gamma}|\}$ that follows a multivariate normal distribution (MVN) with mean 0 and covariance $\sigma^2_{\epsilon}\boldsymbol{V}_{\boldsymbol{\gamma}}$; and $\boldsymbol{V}_{\gamma}$ is the corresponding sub-covariance-matrix of SNPs with $\gamma_i = 1$, $\boldsymbol{V}_{p\times p} = diag(\sigma^2_{q,1}, \cdots, \sigma^2_{q,p})$, where $\sigma^2_{q,i} = \sigma^2_{cis}$ if the $i$th SNP is of cis- annotation and $\sigma^2_{q,i} = \sigma^2_{trans}$ if the $i$th SNP is of trans- annotation. The expectation of the latent indicator variable $(E[\gamma_i])$ is the posterior probability $(PP_i)$ for the $i$th SNP to be an eQTL with effect size $w_i$.

##### 3.1.3 Bayesian Inference

From the above assumed BVSR model (2, 3, 4), the posterior joint distribution of $(\boldsymbol{w}, \gamma, \boldsymbol{\sigma^2}, \boldsymbol{\pi}, \tau)$ is proportional to the product of likelihood and prior density functions,

$$P(\boldsymbol{w}, \gamma, \boldsymbol{\sigma^2}, \boldsymbol{\pi}, \sigma_\epsilon^2 | \mathcal{E}_g, \boldsymbol{X}, \boldsymbol{A}) \propto P(\mathcal{E}_g | \boldsymbol{X}, \boldsymbol{w}, \gamma, \sigma_\epsilon^2) P(\boldsymbol{w} | \boldsymbol{A}, \boldsymbol{\pi}, \boldsymbol{\sigma^2}, \sigma_\epsilon^2) P(\gamma | \boldsymbol{\pi}) \\ P(\boldsymbol{\pi}) P(\boldsymbol{\sigma^2}) P(\sigma_\epsilon^2), \tag{5}$$

where $\boldsymbol{\pi} = (\pi_{cis}, \pi_{trans})$, $\boldsymbol{\sigma^2} = (\sigma_{cis}^2, \sigma_{trans}^2)$, and $\boldsymbol{A}$ is a $p \times 2$ matrix with binary values denoting if the analyzed SNPs are categorized as cis- or trans-.

The main challenge for modeling trans-SNPs in addition of cis-SNPs in the regression model of expression quantitative traits (2, 3, 4) is the computation burden of making Bayesian inference for $(\boldsymbol{w}, E[\gamma] = \boldsymbol{PP})$ with respect to genome-wide genes ($\sim$20K). In order to make the Bayesian inference feasible in practice, we utilize the scalable Expectation-Maximization Markov chain Monte Carlo (EM-MCMC) algorithm proposed for BFGWAS [4]. Specifically, we first segment considered genotype data into approximately independent blocks based on the block-wise linkage disequilibrium (LD) structure of the human genome, i.e., $\boldsymbol{X} = \{\boldsymbol{X}_1, \boldsymbol{X}_2, \cdots, \boldsymbol{X}_K\}$. Then we can write the likelihood function (5) as a product of likelihood functions for $\boldsymbol{X}_k$,

$$P(\mathcal{E}_g | \boldsymbol{X}, \boldsymbol{w}, \gamma, \sigma_\epsilon^2) = \prod_{k=1}^{K} P_k(\mathcal{E}_g | \boldsymbol{X}_k, \boldsymbol{w}_k, \gamma_k, \sigma_\epsilon^2), \tag{6}$$

where $(\mathcal{E}_g | \boldsymbol{X}_k, \boldsymbol{w}_k, \gamma_k, \tau) \sim MVN_{|\boldsymbol{\gamma_k}|}(\boldsymbol{X}_k \boldsymbol{w}_k, \sigma_\epsilon^2 \boldsymbol{I}_{|\boldsymbol{\gamma_k}|})$.

For the convenience of implementing MCMC algorithm per genome block in parallel with given shared hyper parameters $\boldsymbol{\pi} = (\pi_{cis}, \pi_{trans})$, $\boldsymbol{\sigma^2} = (\sigma_{cis}^2, \sigma_{trans}^2)$, $\sigma_\epsilon^2$ is fixed across all genome blocks as the variance of $\mathcal{E}_g$. As shown by previous studies [4], this assumption only results in slightly conserved estimates for $(\boldsymbol{w}, \boldsymbol{PP})$ but saves the hassle of estimating a block-specific error variance. By EM-MCMC algorithm, we estimate $(\boldsymbol{w}_k, \boldsymbol{PP}_k)$ by implementing MCMC algorithm [5, 6] per block (i.e., Expectation step (E-step)) with given values of $(\boldsymbol{\pi}, \boldsymbol{\sigma^2})$; and then update the estimates of $(\boldsymbol{\pi}, \boldsymbol{\sigma^2})$ by maximizing the corresponding expected posterior likelihood function [7] (Maximization step (M-step)) given the estimates of $(\boldsymbol{w}, \boldsymbol{PP})$ from the previous E-step. A few such EM iterations will be run until the estimates of $(\boldsymbol{\pi}, \boldsymbol{\sigma^2})$ converge, which generally requires $\sim$5 EM iterations [4]. The estimates $(\widehat{\boldsymbol{w}}, \widehat{\boldsymbol{PP}})$ from the last E-step will be used to calculate Bayesian genetically regulated gene expression (GReX) levels for additional samples with GWAS genotype data $\widetilde{\boldsymbol{X}}$,

$$\widehat{GReX} = \sum_{i=1}^{p} \widetilde{X}_i (\widehat{PP}_i \widehat{w}_i). \tag{7}$$

Detailed derivations about the conditional posterior distributions of $\boldsymbol{w}_k$ and $\boldsymbol{\gamma_k}$ as well as the EM-MCMC algorithm are referred to the supplementary note of the BFGWAS paper [4]. To further reduce the computation time for fitting such tissue-gene-specific Bayesian model for genome-wide genes, we propose a novel computational technique to implement the MCMC algorithm using only the pre-calculated LD coefficients and summary statistics from standard eQTL analyses based on single variant tests. Below, we briefly outline steps of the EM-MCMC algorithm using only summary statistics.

##### 3.1.4 Fast MCMC using Only Summary Statistics

As shown in the BFGWAS supplementary note [4], the posterior distribution for $(\boldsymbol{w}_k, \boldsymbol{\gamma}_k)$ of block $k$ is

$$P(\boldsymbol{w}_k, \boldsymbol{\gamma}_k | \boldsymbol{X}_k, \boldsymbol{\mathcal{E}}_g, \boldsymbol{\pi}, \boldsymbol{\sigma^2}, \epsilon_\epsilon^2) \propto P(\boldsymbol{\mathcal{E}}_g | \boldsymbol{X}_k, \boldsymbol{w}_k, \boldsymbol{\gamma}_k, \epsilon_\epsilon^2) P(\boldsymbol{w}_k | \boldsymbol{\gamma}_k, \boldsymbol{\sigma^2}, \epsilon_\epsilon^2) P(\boldsymbol{\gamma}_k | \boldsymbol{\pi}). \quad (8)$$

The conditional posterior distribution of $\boldsymbol{w}_{|\boldsymbol{\gamma_k}|}$ is given by

$$\begin{aligned} & P(\boldsymbol{w}_{|\boldsymbol{\gamma_k}|} | \boldsymbol{X}_{|\boldsymbol{\gamma_k}|}, \mathcal{E}_g, \boldsymbol{\gamma}_k, \boldsymbol{\sigma^2}, \sigma_\epsilon^2) \sim \\ & \qquad MVN_{|\gamma_k|} \left( (\boldsymbol{X}'_{|\boldsymbol{\gamma_k}|} \boldsymbol{X}_{|\boldsymbol{\gamma_k}|} + \boldsymbol{V}^{-1}_{|\boldsymbol{\gamma_k}|})^{-1} (\boldsymbol{X}'_{|\boldsymbol{\gamma_k}|} \mathcal{E}_g),\ \sigma_\epsilon^2 (\boldsymbol{X}'_{|\boldsymbol{\gamma_k}|} \boldsymbol{X}_{|\boldsymbol{\gamma_k}|} + \boldsymbol{V}^{-1}_{|\boldsymbol{\gamma_k}|})^{-1} \right). \end{aligned} \quad (9)$$

After integrating $\boldsymbol{w}_k$ out from (8), the marginal conditional posterior distribution for $\boldsymbol{\gamma}_k$ is given by

$$\begin{aligned} P(\boldsymbol{\gamma}_k | \boldsymbol{X}_k, \boldsymbol{\mathcal{E}}_g, \boldsymbol{\pi}, \boldsymbol{\sigma^2}, \sigma_\epsilon^2) &\propto \int_{\boldsymbol{w}_k} P_k(\boldsymbol{\mathcal{E}}_g | \boldsymbol{X}_k, \boldsymbol{w}_k, \boldsymbol{\gamma}_k, \sigma_\epsilon^2) P(\boldsymbol{w}_k | \boldsymbol{\gamma}_k, \boldsymbol{\sigma^2}, \sigma_\epsilon^2) P(\boldsymbol{\gamma}_k | \boldsymbol{\pi}) d\boldsymbol{w}_k \\ &\propto |\boldsymbol{\Omega}_{|\boldsymbol{\gamma_k}|}|^{-1/2} \exp\left\{ \frac{1}{2\sigma_\epsilon^2} (\boldsymbol{\mathcal{E}}'_g \boldsymbol{X}_{|\boldsymbol{\gamma_k}|}) \boldsymbol{V}_{|\boldsymbol{\gamma_k}|} \boldsymbol{\Omega}^{-1}_{|\boldsymbol{\gamma_k}|} (\boldsymbol{X}'_{|\boldsymbol{\gamma_k}|} \boldsymbol{\mathcal{E}}_g) \right\} P(\boldsymbol{\gamma}_k | \boldsymbol{\pi}), \end{aligned} \quad (10)$$

where $\boldsymbol{\Omega}_{|\boldsymbol{\gamma_k}|} = \boldsymbol{V}_{|\boldsymbol{\gamma_k}|} (\boldsymbol{X}'_{|\boldsymbol{\gamma_k}|} \boldsymbol{X}_{|\boldsymbol{\gamma_k}|}) + \boldsymbol{I}_{|\boldsymbol{\gamma_k}|}$, $\sigma_\epsilon^2$ will be taken as the variance of $\boldsymbol{\mathcal{E}}_g$, and the subscript $|\gamma_k|$ indicates sub-matrices or sub-vectors corresponding to variants with nonzero indicator variables. That is, $\boldsymbol{V}_{|\boldsymbol{\gamma_k}|}$ is a diagonal matrix with $(\boldsymbol{V}_{|\boldsymbol{\gamma_k}|})_{jj} = \sigma^2_{cis}$ or $\sigma^2_{trans}$ given the $j$th SNP is cis- or trans-.

As discussed in the BFGWAS paper [4], majority computation time is spent on implementing the MCMC algorithm per genome block (E-step), because $> 10,000$ MCMC iterations are required for obtaining converged estimates for $(\boldsymbol{w_k}, \boldsymbol{\gamma}_k)$. Particularly, most computation resource is spent on evaluating the posterior likelihood (9) and (10) per MCMC iteration, where calculating $\boldsymbol{X}'_{|\boldsymbol{\gamma_k}|} \boldsymbol{X}_{|\boldsymbol{\gamma_k}|}$ and $\boldsymbol{X}'_{|\boldsymbol{\gamma_k}|} \mathcal{E}_g$ costs most computation time.

Therefore, by deriving values of $\boldsymbol{X}'_{|\gamma_k|}\boldsymbol{X}_{|\gamma_k|}$ and $\boldsymbol{X}'_{|\gamma_k|}\mathcal{E}_g$ from pre-calculated LD coefficients and summary statistics [8], up to $90\%$ computation time can be saved.

Consider the single variant regression model that is generally used for standard eQTL analyses,

$$\mathcal{E}_g = X_i w_i + \boldsymbol{\epsilon}, \tag{11}$$

where $X_i$ denotes the genotype vector of the $i$th SNP. The least square estimator $\widetilde{w}_i$ is given by $\widetilde{w}_i = (X_i'X_i)^{-1}X_i'\mathcal{E}_g$. Then the elements of vector $\boldsymbol{X}'_{|\gamma_k|}\mathcal{E}_g$ are given by

$$(\boldsymbol{X}'_{|\gamma_k|}\mathcal{E}_g)_i = \widetilde{w}_i(X_i'X_i), \tag{12}$$

where $X_i'X_i = (n-1)Var(X_i)$ can be either pre-calculated using individual-level genotype data or approximated by $2nf_i(1-f_i)$ with sample size $n$ and minor allele frequency $f_i$.

Note that $\{X_i'X_i;\ i = 1, ..., p\}$ are also the diagonal values of $\boldsymbol{X}'_{|\gamma_k|}\boldsymbol{X}_{|\gamma_k|}$. For the off-diagonal values, $[\boldsymbol{X}'_{|\gamma_k|}\boldsymbol{X}_{|\gamma_k|}]_{(i,j)}$ can be derived from the LD coefficient between the $i$th and $j$th SNPs, $r_{ij} = \frac{X_i'X_j}{\sqrt{(X_i'X_i)(X_j'X_j)}}$. That is,

$$[\boldsymbol{X}'_{|\gamma_k|}\boldsymbol{X}_{|\gamma_k|}]_{ij} = r_{ij}\left(\sqrt{(X_i'X_i)(X_j'X_j)}\right). \tag{13}$$

Thus, given the summary statistics of the variance of quantitative gene expression trait $\mathcal{E}_g$, sample size $n$, either SNP genotype variances $\{Var(X_i)\}$ or minor allele frequencies $\{f_i\}$, pre-calculated LD coefficients $\{r_{i,j};\, i,j = 1,\cdots,p\}$, and least square estimates of effect sizes $\{\widetilde{w}_i\}$ from single variant regression models (12), we can obtain values of $\boldsymbol{X}'_{|\gamma_k|}\boldsymbol{X}_{|\gamma_k|}$ and $\boldsymbol{X}'_{|\gamma_k|}\mathcal{E}_g$ with manageable computation cost to evaluate the posterior mean of $\boldsymbol{w}_k$ (9) and the posterior likelihood for $\boldsymbol{\gamma}_k$ (10). Moreover, if individual level genotype and expression quantitative trait data are not available, the MCMC can still be implemented by using summary statistics where minor allele frequencies and LD coefficients could be approximated by corresponding values generated from reference panels of the same ethnicity.

##### 3.1.5 Adapted EM-MCMC Algorithm

To further reduce computation burden, instead of considering all segmented genome blocks for genome-wide SNPs, we only fit model (2) with pruned genome blocks that either contain at least one cis-SNP or at least one potential trans-eQTL with p-value $< 1 \times 10^{-5}$ by single variant test (i.e., testing $H_0 : w_i = 0$ with model (12)).

The steps of our adapted EM-MCMC algorithm per gene per tissue type are as follows:

(i) Generate summary statistics: either obtain from individual level genotype data and single variant analyses for genome-wide SNPs, or obtain SNP genotype variances and LD coefficients from reference panel and other summary statistics from previous standard eQTL analyses based on single variant tests;

(ii) Prune genome blocks for applying model (2):

   (a) Consider blocks that contain either at least one cis-SNP or at least one potential trans-eQTL with single variant test p-value $< 1 \times 10^{-5}$;

   (b) Select up to $B$ blocks with minimal $p$-values within block from smallest to largest. For example, $B = 100$ was used in our application studies. This number can be tuned by users to reduce total computation time accordingly;

   (c) Select any remaining blocks containing cis-SNPs that were not selected in (b);

(iii) Apply EM-MCMC algorithm to the pruned blocks:

   (a) Fix $\sigma_\epsilon^2$ at the variance of $\mathcal{E}_g$;

   (b) Set initial values for $(\boldsymbol{\pi}, \boldsymbol{\sigma^2})$, e.g., $\pi_{cis} = \pi_{trans} = 1 \times 10^{-6}$ and $\sigma_{cis}^2 = \sigma_{trans}^2 = 0.1$;

   (c) E-step: Conditioning on the most recent estimates of $(\boldsymbol{\pi}, \boldsymbol{\sigma^2})$, estimate $(\boldsymbol{w}, \boldsymbol{PP})$ by implementing MCMC algorithm per block using only summary statistics;

   (d) M-step: Conditioning on the estimates of $(\boldsymbol{w}, \boldsymbol{PP})$ from the previous E-step, update $(\boldsymbol{\pi}, \boldsymbol{\sigma^2})$ by their maximum a posteriori estimates (MAPs), maximizing the expected log-posterior-likelihood functions [7];

   (e) Repeat the EM-steps (c) and (d) for a few times until the MAPs of $(\boldsymbol{\pi}, \boldsymbol{\sigma^2})$ converge, e.g., 5 EM steps.

(iv) Estimates of $(\boldsymbol{w}, \boldsymbol{PP})$ from the last E-step will be used to impute Bayesian GReX from GWAS genotype data of new samples by (7).

See the supplementary note of the BFGWAS paper [4] for details of the EM-MCMC algorithm (iii).

##### 3.2 Software

Software implementing this BGW-TWAS is now available at GitHub (`https://github.com/yanglab-emory/BGW-TWAS`). All steps are wrapped together (enabling parallel computation) through submitting jobs by a Makefile that is generated by a Perl script, which wraps together generating summary statistics by single variant tests with individual-level genotype and gene expression data, pruning genome blocks, implementing adapted EM-MCMC algorithm, and calculating Bayesian $\widehat{GReX}$ for TWAS.

#### 3.3 ROS/MAP Data Description

##### 3.3.1 Study Design

ROS and MAP are prospective cohort studies of aging and dementia, which recruit older adults without known dementia at enrollment who agree to annual clinical testing and brain donation with structured collection of postmortem brain indices at the time of death. All participants sign an informed consent, an Anatomic Gift Act and a consent for their data deposited in the Rush Alzheimer's Disease Center (RADC) repository to be re-purposed by other investigators. Both studies were approved by an Institutional Review Board of Rush University Medical Center, Chicago, IL. Both studies employ harmonized clinical and postmortem data collection facilitating joint analyses.

##### 3.3.2 Cognitive testing and Cognitive Status Diagnosis

Trained technicians administered 17 cognitive tests annually as described previously from which a composite measure of global cognition was constructed[9]. Cognitive status was determined in a three-step process. Annual cognitive testing was scored by a computer and reviewed by a neuro-psychologist to diagnose cognitive impairment and reviewed by a physician. At the time of death, a physician used all cognitive and clinical data collected prior to death, blinded to all postmortem data, to classify persons with respect to dementia, mild cognitive impairment and no cognitive impairment as previously described[10].

#### 3.4 Postmortem Assessment

Brain removal, tissue sectioning and preservation, and a uniform gross and microscopic examination with quantification of post-mortem indices followed a standard protocol [11, 12, 13]. AD pathology (i.e., neuritic plaques, diffuse plaques and neurofibrillary tangles) was visualized using a modified Bielschowsky silver stain on sections from five brain regions and a global summary measure was constructed as described in prior publications[14]. $\beta$-amyloid load and paired helical filament tau immunoreactive neuronal neurofibrillary tangles (tangles) were quantified in 8 brain regions (anterior cingulate cortex, superior frontal cortex, mid frontal cortex, inferior temporal cortex, hippocampus, entorhinal cortex, angular gyrus/supramarginal gyrus, and calcarine cortex). Overall $\beta$-amyloid load was calculated through averaging mean percent area of $\beta$-amyloid deposition per region, across multiple brain regions. Likewise, tangles densities were derived by averaging tangles densities across corresponding brain regions. The global measure of AD pathology is based on counts of neuritic and diffuse plaques and neurofibrillary tangles (15 counts) on 6m sections stained with modified Bielschowsky.
